# γ-secretase promotes postsynaptic maturation through the cleavage of a Wnt receptor

**DOI:** 10.1101/2020.11.18.387720

**Authors:** Lucas Restrepo, Alison DePew, Elizabeth Moese, Stephen Tymanskyj, Michael Parisi, Michael Aimino, Juan Carlos Duhart, Hong Fei, Timothy J. Mosca

## Abstract

An emerging feature of neurodegenerative disease is synaptic dysfunction and loss, leading to the suggestion that mechanisms required for synaptic maturation may be linked to disease. Synaptic maturation requires the transmission of signals between nascent synaptic sites and the nucleus, but how these signals are generated is not well understood. We posit that proteolytic cleavage of receptors, which enables their translocation to the nucleus, may be a shared molecular mechanism between the events that promote synaptic maturation and those linked to later-onset disorders of the nervous system, including neurodegenerative disease. Here we show during synaptic development, that cleavage of synaptic maturation molecules requires γ-secretase, a protein complex linked to Alzheimer’s Disease, a devastating neurodegenerative condition, is required for postsynaptic maturation. In the absence of γ-secretase, *Drosophila* neuromuscular synapses fail to appropriately recruit postsynaptic scaffolding and cytoskeletal proteins, and mutant larvae display behavioral deficits. At the NMJ, γ-secretase promotes synaptic maturation through the cleavage of the Wnt receptor Fz2, and the subsequent entry of its C-terminus into the nucleus. A developmental synaptic role for γ-secretase is also conserved in both the *Drosophila* central nervous system and mammalian cortical neuron dendrites. Finally, we found that similar maturation defects are evident in fly models for ALS, Alzheimer’s, Huntington’s, and Parkinson’s Diseases. The previously unknown, but conserved, role for γ-secretase coupled with its well-known role in neurodegenerative disease suggest that neurodevelopmental defects may be common to diverse neurodegenerative disease models.

## Introduction

Postsynaptic development is essential for the transition of neuronal connections from nascent contacts to fully formed synapses that drive robust, reliable neurotransmission. Maturation includes the recruitment of specialized pre- and postsynaptic scaffolding, cytoskeletal, and neurotransmitter release proteins to the synapse (*1–4*) as well as functional refinement including synaptic pruning and activity-dependent maturation (*5–8*). The molecular mechanisms of maturation are incompletely understood but important, as their failure can lead to neurodevelopmental disorders (*9*) and possibly later onset neurodegenerative diseases (*10*).

At the *Drosophila* neuromuscular junction (Fig. 1A), synaptic boutons form from outgrowth of the presynaptic motoneuron (*11*) in an activity-dependent manner (*12, 13*). These boutons initially consist of neuronal membrane and synaptic vesicles (*14*). As the animal develops, these boutons subsequently mature presynaptically by assembling active zones (*15*) and postsynaptically by recruiting neurotransmitter receptors, scaffolding, and cytoskeletal proteins (*16–19*). Failures in the maturation process result in “ghost” boutons, which are discrete synaptic terminals where the presynaptic bouton fails to recruit postsynaptic components like the scaffolding protein Discs Large (*13, 14, 16, 20*). Ghost boutons are comprised of neuronal membrane and synaptic vesicles but lack active zones and any postsynaptic components. Failures in maturation also result in a generalized decrease and thinning of the postsynaptic spectrin cytoskeleton (*16*) at all boutons. Under normal circumstances, maturation failures that manifest as ghost boutons occur infrequently. Mutations that perturb NMJ maturation and development, however, increase ghost bouton prevalence 3- to 20-fold (*13, 16, 21, 22*). One critical molecular mechanism thought to promote such NMJ synaptic bouton maturation (*14, 16, 20*) is the Frizzled2 Nuclear Import (FNI) pathway (Fig. S1A), which involves non-canonical Wnt signaling including the cleavage of the C-terminus of the Wnt receptor Frizzled2 (Fz2-C) and its import into muscle nuclei to influence postsynaptic development (*21, 23*). However, the hypothesis that FNI underlies synaptic maturation has remained controversial and the physiological relevance of the FNI pathway unclear due to potentially confounding overlap with canonical Wnt signaling (*23–25*). Further, and most importantly, it remains unknown what proteases function to promote Fz2 cleavage, which has precluded a direct study of the mechanism of receptor cleavage to understand how proteolysis could gate synaptic maturation. Combined, these three issues have precluded a thorough understanding of the molecular gatekeepers of FNI-dependent synaptic maturation and how critical synaptic mechanisms that underlie neurodevelopment might be conserved across synapses and organisms. To address these three gaps, we set out to determine whether Fz2 cleavage promotes synaptic maturation and identify the relevant protease.

**Fig. 1.**
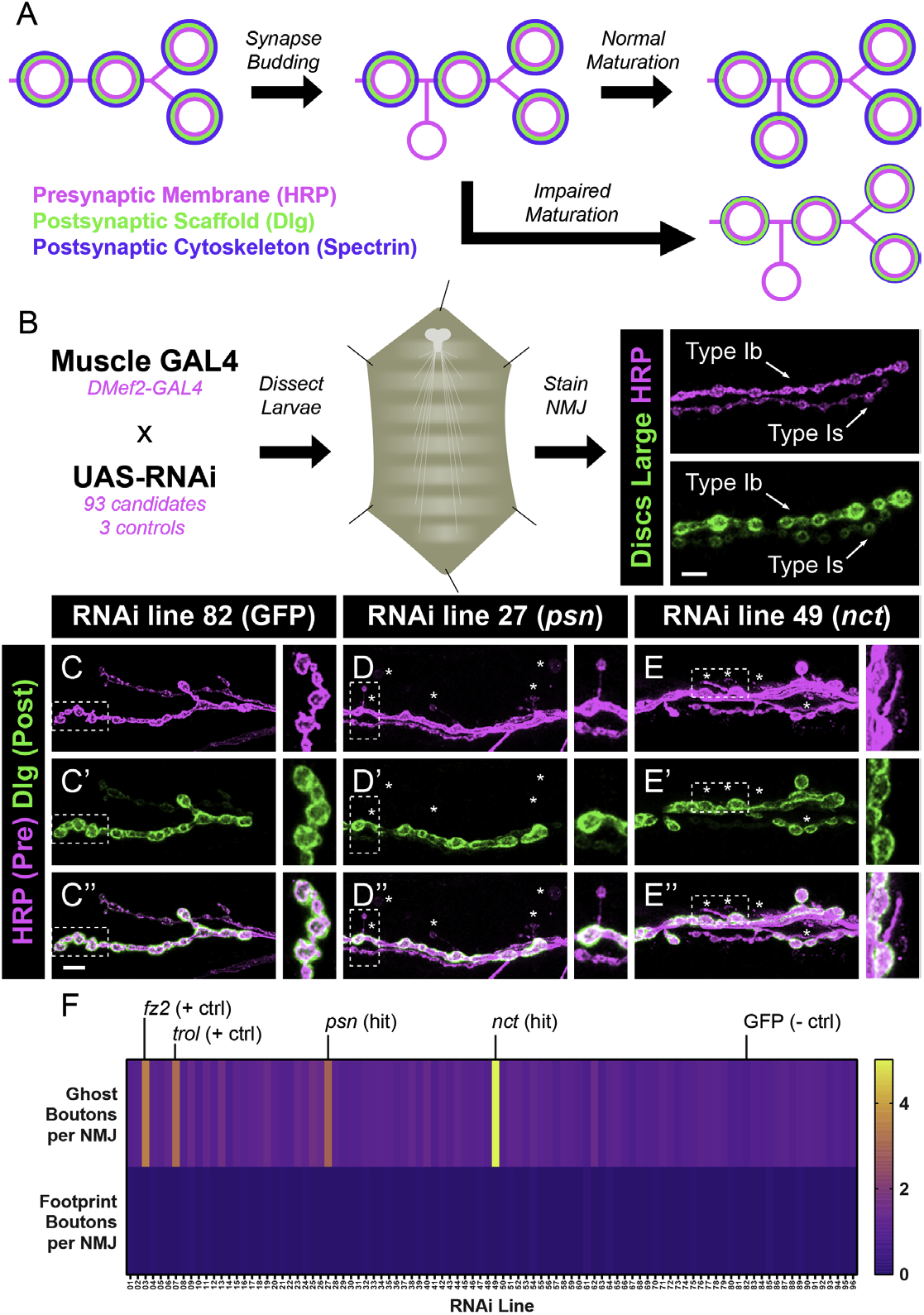
A tissue-specific RNAi screen identifies *presenilin* and *nicastrin* as required for normal postsynaptic development and maturation. (**A**) Schematic of bouton addition at the *Drosophila* neuromuscular junction. Presynaptic boutons (HRP, magenta) are surrounded by the postsynaptic scaffolding protein Discs Large (Dlg, green) and the cytoskeleton (Spectrin, blue). (**B**) Pipeline of the RNAi screen: UAS-RNAi candidates and controls were driven using a muscle-specific GAL4 driver (DMef2-GAL4). Larvae were dissected to the fillet prep and stained with antibodies to Dlg and HRP. The representative confocal micrograph shows that, in a wild-type animal, type Ib and type Is boutons are both positive for HRP and Dlg. (**C – E**) Representative confocal images of NMJ boutons stained with antibodies to HRP (magenta) and Dlg (green) in larvae expressing RNAi against GFP (C), psn (D), or *nct* (E) in muscles. Asterisks indicate ghost boutons. Note the increase over control lines in both psn and nct RNAi. Insets represent individual ghost boutons from the image. Scale bar = 10 μm. Insets represents high magnification ghost or control boutons marked by the dashed line. (**F**) Heat maps of the number of ghost boutons (top) or footprint boutons (bottom) in each gene targeted by RNAi. Four positive hits corresponding to the two positive controls (*fz2* and *trol*), *psn* and nct showed an increase in ghost boutons while the negative control (GFP) showed no phenotype. No lines showed an increase in footprints, suggesting that the manipulations are not associated with destabilization / degeneration.

### An *in vivo* Screen to Identify Proteases Involved in Synaptic Maturation

We first sought to directly test whether Fz2 cleavage promotes synaptic maturation. Though Fz2 cleavage promotes synaptic growth (*23*) and nuclear Fz2-C entry is required for normal maturation (*16*), it is not known if blocking it impairs maturation. In *fz2* null mutants, there is a four-fold increase in ghost boutons (Fig. S1B-C,F) (*16*), indicating that the receptor is indeed required for maturation. To specifically test if the cleavage event is required, we used a molecular replacement approach (Fig. S1E) by expressing Fz2 transgenes in the postsynaptic muscles of *fz2* null mutants that encode the full-length wild-type receptor, a non-cleavable receptor, or a truncated receptor form consisting only of the receptor C-terminus (*23*). We found that muscle expression of full-length Fz2 or just the Fz2 C-terminus rescues the maturation defect, but non-cleavable Fz2 does not (Fig. S1F). The residues that comprise the Fz2 cleavage site (KTLES) and which are mutated in the uncleavable form, also serve as the essential residues that make up the binding site of the canonical Wnt signaling pathway protein Dishevelled (Dsh) (*25*). Our experiment indicated that the KTLES sequence was essential for maturation but the above experiment alone could not rule out that KTLES was functioning as a Dsh binding site (and not a cleavage site) as a potential mechanism to promote maturation. As canonical Wnt signaling functions in postsynaptic muscles and Dsh is present during NMJ development (*24*), we tested whether specifically impairing canonical Wnt signaling through Dsh would perturb maturation. This would allow us to differentiate between the two downstream pathways and determine if the KTLES sequence is functioning as a cleavage site or a Dsh-binding site to promote NMJ maturation. To distinguish between these two possibilities, we expressed a dominant-negative Dsh^DIX^ construct that impairs canonical signaling downstream of Fz2 (*25*) in muscles and measured maturation to test specifically whether postsynaptic Dsh was required for synaptic maturation. Disruption of Dsh in muscle manipulation did not affect ghost bouton number (Fig. S1D,F), suggesting that Dsh is dispensable for NMJ maturation. Thus, the failed maturation seen when the KTLES site is mutated (Fig. S1F) is likely due to impaired Fz2-C cleavage and not as a result of impaired Dsh binding (as Dsh is not needed for maturation). These results highlight the requirement for Fz2 cleavage, and with it, the Frizzled2 Nuclear Import pathway, as a critical mediator of synaptic maturation.

With the importance of Fz2 cleavage defined, we sought to identify the proteases responsible to understand more about the mechanisms that serve as gatekeepers of synaptic maturation. As the Fz2 cleavage site is a consensus glutamyl endopeptidase site (*23*), we performed a tissue-specific RNAi screen (*26*) against 93 candidates predicted by GO-term analysis to have glutamyl endopeptidase activity (Fig. 1B), to identify proteases that promote Fz2 cleavage. We reasoned that muscle-specific RNAi would 1) target the appropriate tissue as Fz2 is expressed in muscle and 2) positive hits would cause similar maturation phenotypes as the loss of Fz2 itself. We dissected wandering third instar larvae and stained neuromuscular junctions (NMJs) for pre-and postsynaptic markers (HRP and Dlg) and quantified ghost boutons in both type Ib (big) and Is (small) terminals at the muscle 6/7 NMJ (*12, 13, 16*). We also quantified “footprint” boutons, which occur when the presynaptic NMJ degenerates, leaving postsynaptic Dlg staining behind (*27–29*); these can indicate synaptic degeneration and serve as a control to distinguish developmental defects from destabilization. On average, control animals have only 1 ghost bouton at the muscle 6/7 NMJ (*14, 16, 20, 30, 31*) – a positive screen hit was denoted by a statistically significant increase in ghost bouton number. In negative control larvae with RNAi against an innocuous GFP transgene (Fig. 1C), the incidence of ghost boutons was not increased. In positive control larvae expressing RNAi against the *dfz2* receptor itself (*23*) or the secreted protein *trol* which is required for Wnt signaling at the NMJ (*20*), maturation was severely impaired, resulting in a 3-fold increase in ghost boutons (Fig. 1F). Only two other RNAi lines (Fig. 1D-E) significantly increased ghost bouton number, those targeting *presenilin* and *nicastrin*. These errors were not accompanied by footprint phenotypes (Fig. 1F), suggesting that *presenilin* and *nicastrin* RNAi specifically impaired development.

Presenilin (Psn) and Nicastrin (Nct), along with Aph-1 and Pen-2, comprise γ-secretase, a proteolytic holocomplex whose function is linked to the devastating neurodegenerative condition known as Alzheimer’s Disease (*32, 33*). Mutations in *PSEN1*, the human Presenilin homologue, are the most widely-known genetic cause of familial early-onset Alzheimer’s Disease (AD) (*34–36*). Presenilin is the catalytic subunit of γ-secretase and these mutations alter the cleavage of the Amyloid Precursor Protein (APP), which is at least partially responsible for AD pathology (*35*), though this has remained controversial (*37*). Despite clear clinical relevance, the biological functions of Presenilin and γ-secretase remain incompletely understood among roles in cell-cell signaling (*33*), axon guidance (*38*), vesicle cycling (*39, 40*), and neuronal development (*41–43*). Our identification of Psn and Nct in a genetic screen raised the intriguing possibility that γ-secretase could promote postsynaptic maturation via the mechanism of Fz2 cleavage.

### Muscle γ-secretase Promotes Postsynaptic Maturation and Normal Behavior

We first examined whether γ-secretase localizes endogenously to NMJ synapses. We detected both Presenilin (Fig. 2A) and Nicastrin (Fig. 2C) that overlapped with and extended beyond presynaptic staining. This extension overlapped with postsynaptic Dlg staining, suggesting that Presenilin and Nicastrin localize both pre- and postsynaptically, forming two pools of γ-secretase at the NMJ. This is consistent with both previous work (*44*) and our screen results. These signals disappear in null mutants of either gene (Fig. 2B, D), demonstrating the specificity of the staining patterns. We also detected that muscle-driven, epitope-tagged Presenilin and Nicastrin can localize to NMJs (Fig. S2A-D) and olfactory neuron-driven Presenilin and Nicastrin can localize to CNS synapses (Fig. S2E-F). This data indicates that γ-secretase components are localized to both central and peripheral synapses, and specifically at NMJ terminals, comprise a postsynaptic pool.

**Fig. 2.**
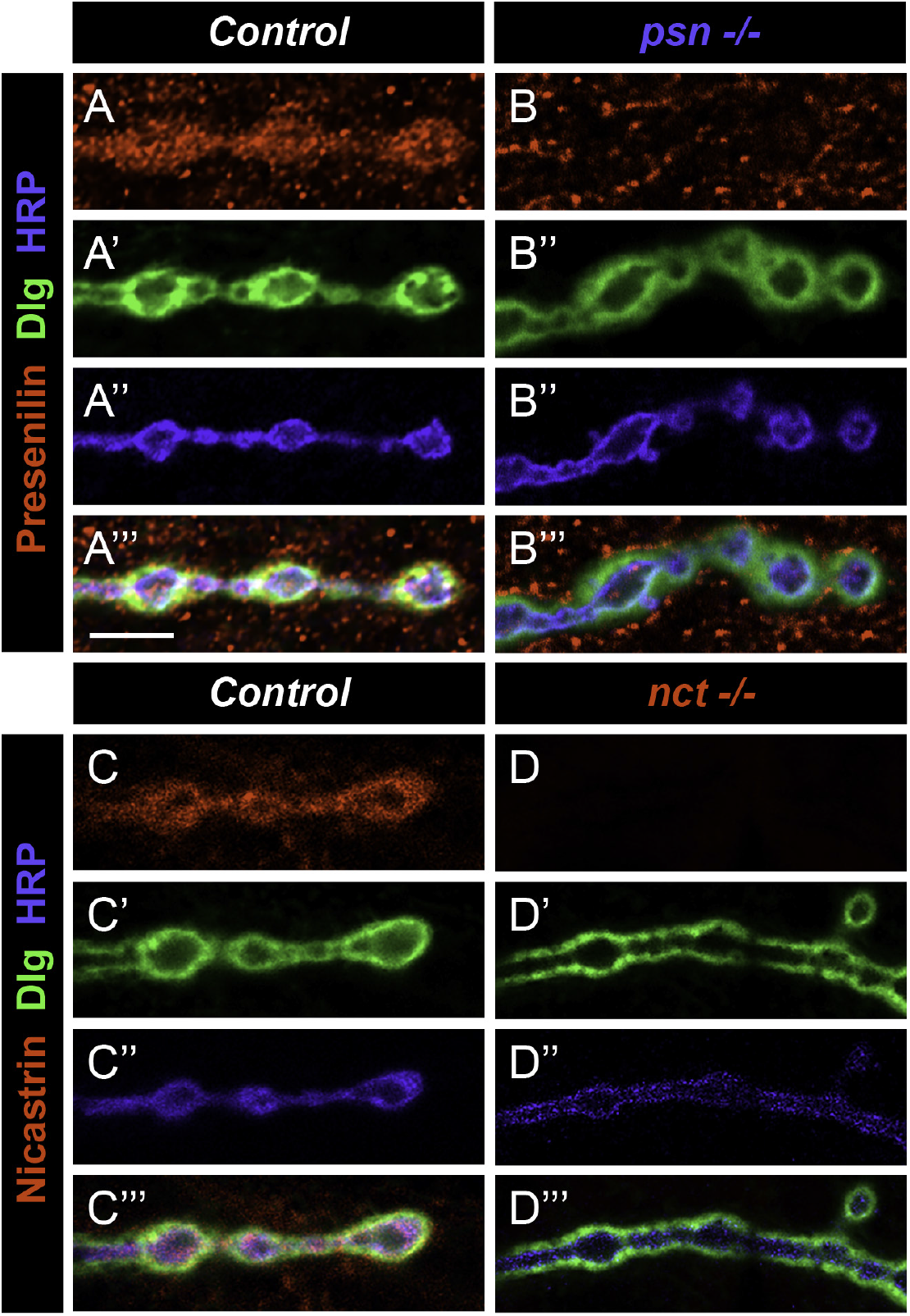
Presenilin and Nicastrin are expressed pre- and postsynaptically at developing NMJs. (**A – B**) Representative single confocal section images taken using Fast AiryScan of control (A) or *psn* null mutant (B) larvae stained with antibodies against Presenilin (red), Dlg (green), and HRP (blue). Psn staining is evident within the HRP-positive bouton, indicating presynaptic localization. Psn staining observed outside the HRP-positive region is within the Dlg-positive region, suggesting an additional postsynaptic pool of staining. Psn staining is absent in *psn* null mutants, demonstrating the specificity of the antibody. (**C – D**) Representative single confocal section images taken using Fast AiryScan of control (C) or *nct* null mutant (D) larvae stained with antibodies against Nicastrin (red), Dlg (green), and HRP (blue). A similar pattern is evident: Nicastrin staining is both coincident with presynaptic HRP staining and postsynaptic Dlg staining, indicating both pre- and postsynaptic pools of Nicastrin. Nct staining is absent in *nct* null mutants, indicating antibody specificity. Scale bar = 5 μm.

We next sought to determine whether the loss of γ-secretase components Presenilin and Nicastrin (Fig. 3A) resulted in failures in synaptic maturation. We examined null *psn* and *nct* mutants and quantified ghost bouton number and postsynaptic α-spectrin staining intensity. Each mutant displayed a significant increase in the number of ghost boutons (Fig. 3C-D) and a marked reduction in postsynaptic α-spectrin (Fig. 3G-H) over controls (Fig. 3B,F), demonstrating impaired maturation. We observed similar defects in mutants for the other two subunits of the γ-secretase complex, Aph-1 and Pen-2 (Fig. S3). These defects were independent of other synaptic defects, as bouton number, muscle area, active zone and glutamate receptor density, and synaptic vesicle protein levels were all unchanged in these mutants (Fig. S4), suggesting the defects were specific to maturation and not secondary to other morphological faults. To subsequently assess if these phenotypes were due to the enzymatic function of γ-secretase or some other role, we examined whether pharmacological blockade of γ-secretase activity could impair synaptic maturation at the *Drosophila* NMJ. When wild-type flies were fed the potent γ-secretase inhibitor L645,458 (*43*) in standard *Drosophila* media, they displayed a 5-fold increase in ghost boutons (Fig. S5A-C). Importantly, L645,458 did not enhance the morphological maturation phenotype of a *psn* null mutant (Fig. S5C), suggesting that the phenotype observed in that mutant is due predominantly to the activity of γ-secretase.

**Fig. 3.**
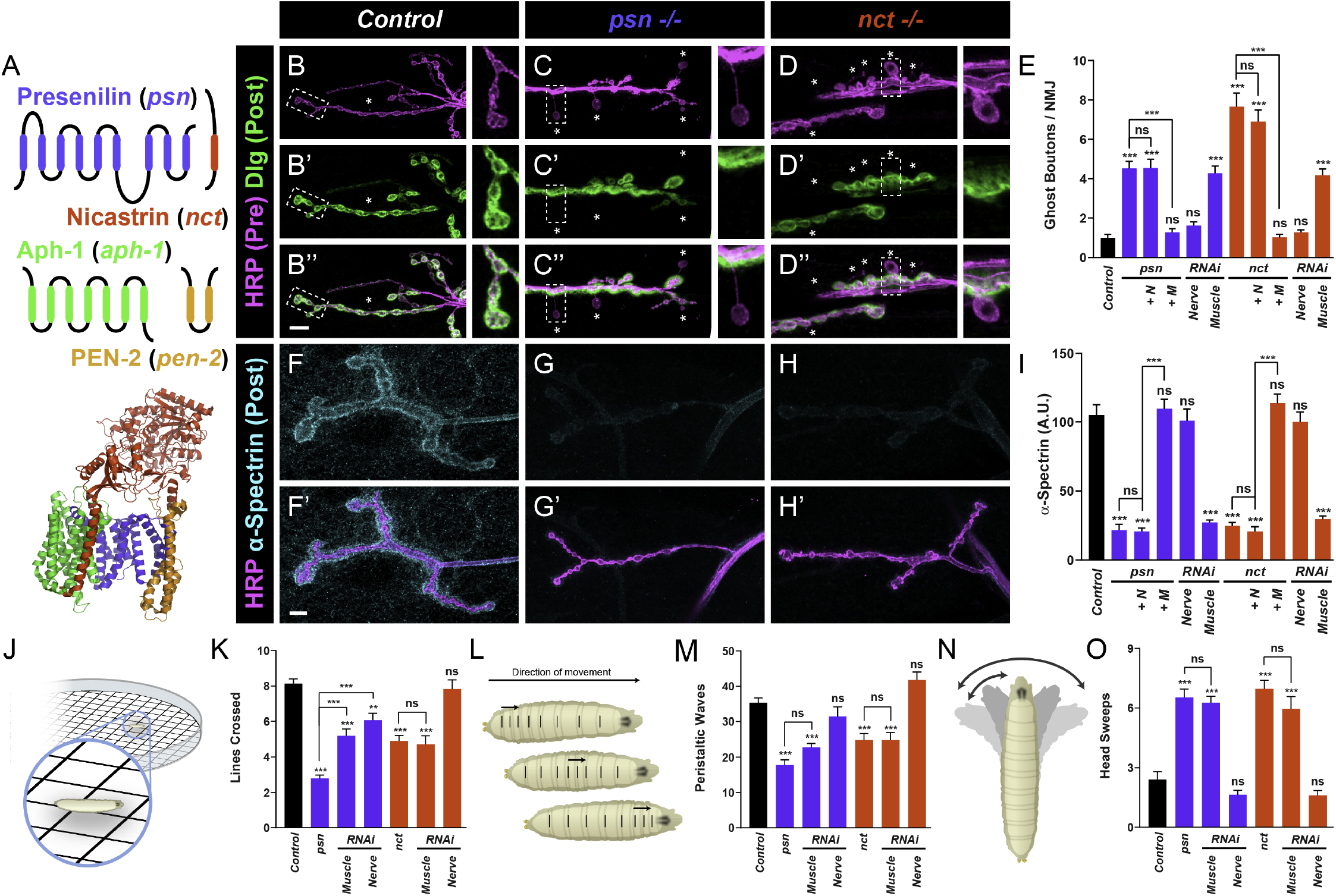
Loss of postsynaptic *psn* or *nct* impairs postsynaptic development and function. (**A**) Schematic of individual γ-secretase subunits (top): Presenilin (blue), Nicastrin (red), Aph-1 (green), and Pen-2 (orange) and ribbon diagram (bottom) representing the structure of the γ-secretase holocomplex (*79*). (**B – D**) Representative confocal images of control (B), *psn* mutant (C), and *nct* mutant (D) larvae stained with antibodies against HRP (magenta) and Dlg (green). Asterisks indicate ghost boutons; insets represent high magnification ghost or control boutons marked by the dashed line. (**E**) Quantification of ghost boutons in mutant, rescue, and RNAi genotypes. *psn* and *nct* mutants show a 5- to 7-fold increase in ghost boutons over control genotypes. The ghost bouton phenotype can be suppressed by postsynaptic muscle expression of a Psn or Nct transgene in the respective mutant (+ M) but not presynaptic neuronal expression (+ N). Further, the phenotype can be recapitulated by muscle RNAi against *psn* or *nct* but not neuronal RNAi. (**F – G**) Representative confocal images of control (F), *psn* mutant (G), and *nct* mutant (H) larvae stained with antibodies against HRP (magenta) and α-spectrin (cyan). α-spectrin fluorescence and thickness is markedly reduced (but not eliminated) in *psn* and *nct* mutants while HRP staining is unaffected. (**I**) Quantification of α-spectrin fluorescence intensity in mutant, rescue, and RNAi genotypes. α-spectrin is reduced by 80% in both *psn* and *nct* mutants; the α-spectrin phenotype is suppressed by muscle expression (+ M) but not neuronal expression (+ N). Similarly, muscle RNAi but not neuronal RNAi induces a comparable phenotype, further suggesting a postsynaptic role for Psn and Nct. (**J**) Diagram of larval crawling assay to measure motility. (**K**) Quantification of larval motility in mutant and RNAi genotypes. Loss of *psn* and *nct* impairs the number of lines crossed. In the *psn* mutant, the crawling defect is recapitulated in both nerve and muscle RNAi, suggesting that both pre- and postsynaptic pools of Psn are required for normal behavior, consistent with previous work (*44*). In the *nct* mutant, however, the mutant deficit is completely recapitulated by muscle RNAi and not neuronal. (**L**) Diagram of larval peristaltic waves with arrows indicating direction of movement and lines on the larva denoting body wall segments. (**M**) Quantification of peristaltic waves in mutant and RNAi genotypes. Loss of *psn* or *nct* causes a 50% reduction in peristaltic waves. The peristalsis defect is completely recapitulated in muscle RNAi of either gene but not by neuronal RNAi. (**N**) Diagram of the larval head sweep. Arrows and shading indicate directions of motion during the head sweep behavior. (**O**) Quantification of head sweeps in mutant and RNAi behavior. Both *psn* and *nct* mutants show an improper 3-fold increase in head sweeps over control larvae. The behavioral phenotype is recapitulated again by muscle, and not neuronal RNAi. Taken together, the data suggest postsynaptic *psn* and *nct* promote normal postsynaptic morphology, maturation, and larval behavior. For morphological experiments, *n* ≥ 8 larvae, 16 NMJs. For behavioral experiments, *n* ≥ 24 larvae. **, *p* < 0.01, ***, *p* < 0.001, n.s. = not significant. Scale bar = 10 μm.

Given that Presenilin and Nicastrin are expressed pre- and postsynaptically at the NMJ, we sought to determine which pool was responsible for promoting effective synaptic maturation. We used tissue-specific rescue of each mutant with epitope-tagged transgenes (*45*) and found that expression of γ-secretase components in presynaptic neurons of their respective mutant backgrounds could not rescue the maturation defects (Fig. 3E, I). Expression in postsynaptic muscles, however, completely suppressed the respective mutant defects (Fig. 3E, I). The same was true of *aph-1* and *pen-2* mutant rescues (Fig. S3E,I), indicating that all γ-secretase subunits are required postsynaptically for proper morphological synaptic development and maturation.

Impaired synaptic maturation is often accompanied by alterations in synaptic plasticity and coordinated behavior (*46–50*). Consistently, *psn* null mutants have been shown to display impaired synaptic plasticity and basal physiological response (*44*). We wondered if these defects were accompanied by altered behavior, and whether such alterations were shared in mutants for each γ-secretase component. We used coordinated larval crawling as a readout along with mutant analysis and tissue-specific RNAi to assess where each gene functions. *Drosophila* larvae crawl using a peristaltic wave-based motion (*51*), occasionally engaging in head sweeps as a foraging behavior. To assay behavior, we used locomotion assays and quantified distance traveled (Fig. 3J, S3J), the number of peristaltic waves during that distance (Fig. 3L, S3L), and the number of head sweeps during that time (Fig. 3N, S3N). In all cases, loss of γ-secretase components reduced larval motility, both in distance traveled (Fig. 3K, S3K) and the number of peristaltic waves (Fig. 3M, S3M), suggesting impaired locomotion.

Intriguingly, loss of γ-secretase also resulted in more head sweeps (Fig. 3O, S3O). These phenotypes were completely recapitulated by muscle RNAi against γ-secretase while neuronal RNAi showed largely no phenotype. This indicates a clear perturbation of coordinated larval locomotion when postsynaptic γ-secretase is disrupted, demonstrating both morphological and functional consequences to blocking γ-secretase in the muscle.

### γ-secretase Functions in the Fz2 Nuclear Import Pathway to Promote Cleavage

These findings demonstrate clear roles for both Fz2 cleavage and muscle γ-secretase in postsynaptic maturation and normal function. We next sought to determine whether those aspects were mechanistically connected, with the working model that γ-secretase activity is essential for Fz2 cleavage. However, given the established role for γ-secretase in Notch signaling (*33, 52*), we first examined whether perturbation of the Notch pathway impaired maturation. If γ-secretase functioned through Notch, we would expect that perturbing the Notch pathway itself would display similar morphological phenotypes. However, mutations of various components of the Notch pathway including *moleskin, Kuzbanian, appl, mastermind*, and *Notch* itself displayed no defects in synaptic maturation, as measured by the presence of ghost boutons (Fig. S6). Thus, γ-secretase is unlikely to function through the Notch pathway to promote synaptic maturation.

To establish a connection between γ-secretase and the Frizzled2 Nuclear Import (FNI) pathway, we used transheterozygous assays and double mutant analysis to determine if Fz2 and γ-secretase components interacted genetically. Single heterozygotes for *psn, nct*, or *fz2* showed normal synaptic maturation in that there was no increase in ghost boutons over controls (Fig. 4A-B,D). Transheterozygotes between *psn* and *nct, psn* and *fz2*, or *nct* and *fz2*, however, showed a significant increase in ghost boutons at the NMJ (Fig. 4C-D). This interaction was specific for γ-secretase, as transheterozygotes with null alleles for *ten-a*, a transmembrane receptor that impairs maturation via a mechanism independent of FNI (*53*), did not display any genetic interactions (Fig. 4D). Finally, double mutants between *aph-1* and *fz2* did not show an enhancement over either single *aph-1* or *fz2* mutants (compare Fig. 4D and Fig. S3E), further suggesting that γ-secretase and *fz2* function in the same, as opposed to parallel, genetic pathways.

**Fig. 4.**
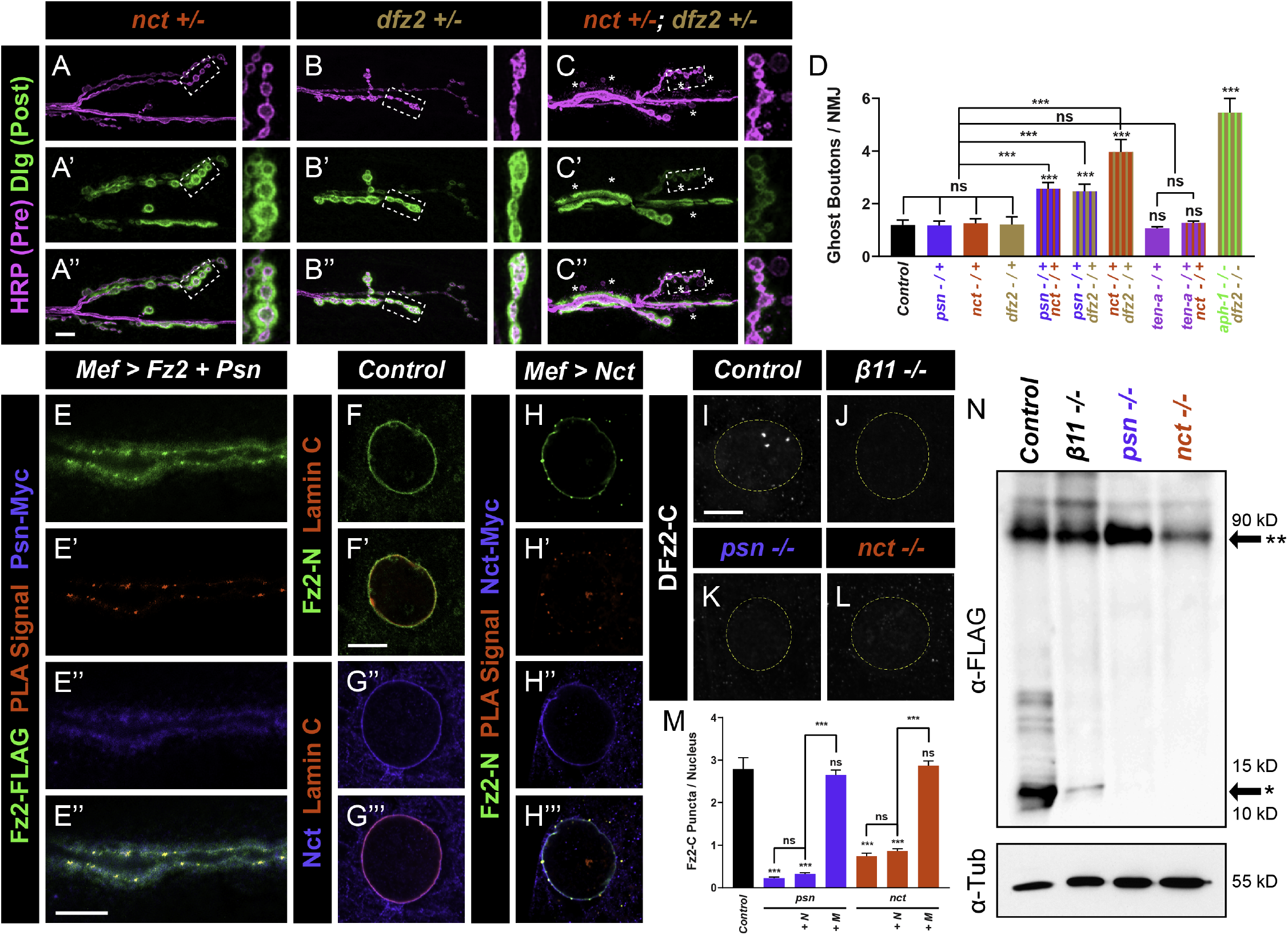
Psn and Nct interact with the Fz2 nuclear import pathway and enable Fz2-C cleavage. (**A – C**) Representative confocal images of *nct* single heterozygotes (A), *dfz2* single heterozygotes (B), and *nct dfz2* double heterozygotes (C) stained for antibodies to HRP (magenta) and Dlg (green). Asterisks indicate ghost boutons. Insets represent high magnification ghost or control boutons marked by the dashed line. Double heterozygotes show an increased incidence of ghost boutons, indicating a genetic interaction. (**D**) Quantification of ghost boutons in genetic interaction genotypes. Single heterozygotes of *psn, nct*, and *dfz2* show no phenotypes, but double heterozygote combinations show increases in ghost boutons, indicating genetic interaction. This interaction is specific for *fz2*, as double heterozygotes with *ten-a*, another mutant that impairs maturation through a different pathway, shows no interaction. Double *aph-1* and *dfz2* null mutants show similar phenotypes to single *aph-1* or *dfz2* mutants. Combined, the data suggest that *γ-secretase* components and *dfz2* function in the same genetic pathway. (**E**) Representative confocal micrograph of larvae expressing Fz2-FLAG and Psn-Myc in postsynaptic muscles, stained for antibodies to FLAG (green) and Myc (blue) and reacted with a proximity ligation assay (red). Both Fz2 and Psn localize postsynaptically and demonstrate positive PLA signal, indicating that they are within 40 nm of each other, a distance consistent with molecular interaction. (**F – G**) Single confocal sections through the center of muscle nuclei in control larvae and stained with antibodies to Fz2-N (F, green) or Nct (G, blue), and Lamin C (red) to mark the nuclear envelope. (**H**) Representative confocal micrograph of larvae expressing Nct-Myc in muscle, stained with antibodies to Fz2-N (green), Myc (blue), and reacted with a proximity ligation assay (red). Positive signal indicates that both Nct and Fz2 are also in close proximity at the nuclear envelope, where Fz2-C cleavage likely occurs (*23*). (**I – L**) Representative confocal images of control (I), *imp-β11* mutant (J), *psn* mutant (K), and *nct* mutant (L) larvae stained with antibodies to Fz2-C. The nuclear border is outlined (dashed lines). All mutants lack nuclear Fz2-C staining, indicating a failure of Fz2-C to enter the nucleus or be cleaved. (**M**) Quantification of Fz2-C puncta in mutant and rescue genotypes. Loss of *psn* or *nct* decreases nuclear Fz2-C puncta by 85-93%. The Fz2-C nuclear defect can be suppressed by muscle expression of the respective transgene but not neuronal expression. This indicates that postsynaptic γ-secretase is required for nuclear Fz2-C entry. (N) Western blot analysis of muscle-expressed Fz2-FLAG cleavage in *control, imp-β11* mutant, *psn* mutant, and *nct* mutant larvae. In all genotypes, the full-length (**) species is observed. In control larvae, the cleaved C-terminus (*) is also observed but is completely absent in *psn* and *nct* mutant larvae, indicating that they are required for Fz2-C cleavage. The Fz2-C cleavage product is observed in *imp-β11* mutant larvae, suggesting that blocking import does not cause cleavage failure as a secondary defect. Its decrease, though, suggests it may be unstable if not imported into the nucleus. Tubulin is used as a loading control. ***, *p* < 0.001. n.s. = not significant. For (A-M), *n* ≥ 6 larvae, 12 NMJs per genotype; (N) is a representative blot of 3 experiments. Scale bars = 10 μm (or 5 μm in E).

Having established γ-secretase and Fz2 in the same genetic pathway, we tested whether they could function together in a complex, consistent with a direct role in promoting cleavage. We examined their synaptic localization and used proximity ligation assays (PLA) (*54*). Epitope-tagged Fz2 and Nct both localize postsynaptically when expressed only in postsynaptic muscles and show positive PLA signal, suggesting they are within close proximity of each other (Fig. 4E). Notably, both Fz2-N (*16, 23, 55*) (Fig. 4F) and Nct (Fig. 4G) localize to the nuclear periphery, where Fz2 cleavage likely occurs (*14, 23*). We also detected positive PLA signal between these pools of Fz2 and Nct (Fig. 4H), indicating that the two are in sufficiently close proximity to mediate a physical interaction.

Both genetic and proximity interactions between Fz2 and γ-secretase suggested their coordinated involvement. If γ-secretase is required for Fz2 cleavage, we hypothesized that this would 1) be biochemically detectable and 2) abrogate nuclear Fz2-C entry (Fig. S1A), as cleavage of the C-terminus is required for nuclear import (*23*). We first assayed nuclear Fz2-C staining, which is indicative of successful Fz2 cleavage and subsequent Fz2-C nuclear import (*14, 16, 21, 23, 30*). Nuclear Fz2-C puncta are evident in control samples (Fig. 4I) and blocking their import results in the absence of nuclear staining (Fig. 4J). We observed a similar loss of nuclear Fz2-C staining in both *psn* and *nct* mutant nuclei (Fig. 4K-L). This loss of nuclear puncta could be suppressed by muscle, but not neuronal, expression of Psn or Nct in the respective mutant background (Fig. 4M), consistent with a role for postsynaptic γ-secretase in promoting nuclear Fz2-C entry, likely acting specifically at the cleavage step. Wingless ligand expression, Fz2 receptor expression and localization as well as trafficking and endocytosis are unchanged by the loss of γ-secretase (Fig. S7) so the defects observed are unlikely to be due to impairing the pathway at an upstream step. To directly assay Fz2 cleavage, we expressed a C-terminally FLAG-tagged Fz2 (*76, 56*) in muscles of control, *psn* or *nct* mutant larvae and examined receptor cleavage in larval body-wall lysates via Western blot. In control samples (Fig.

4N), we observed both full-length FLAG-tagged (90 kDa) and Fz2-C bands (12 kDa). The full-length bands were present in *psn* and *nct* mutants, but the cleaved species was absent, suggesting failed cleavage (Fig. 4N). Moreover, *imp-β11* mutants, which block Fz2-C nuclear entry, still show Fz2-C cleavage (Fig. 4N), suggesting that absent cleavage is not a byproduct of failed nuclear import. Taken together, these indicate that γ-secretase is required for Fz2 cleavage and enables the subsequent nuclear import of Fz2-C.

### Restoring Nuclear Fz2-C Suppresses Maturation Defects in *γ-secretase* Mutants

Our data supports a role for muscle γ-secretase in promoting postsynaptic maturation by enabling Fz2-C cleavage and enabling its nuclear import (Fig. 5A). However, as γ-secretase can cleave many targets (*33*), we sought to determine if Fz2-C nuclear entry was the causative event underlying failed maturation in γ-secretase mutants (Fig. 5A’). If so, we reasoned that restoration of nuclear Fz2-C to γ-secretase mutant muscles (Fig. 5A’’) would suppress the maturation defects. To that end, we expressed a myc-tagged Fz2-C peptide (*16, 20, 23*) with a nuclear localization signal (myc-NLS-Fz2-C) in muscles of *psn* or *nct* mutants and assayed the morphological and larval locomotion phenotypes associated with these mutants. In all cases, muscle Fz2-C expression suppressed the ghost bouton defects (Fig. 5B-F, L), α-spectrin defects (Fig. 5G-K, M), and behavioral abnormalities (Fig. 5N-P) associated with *psn* and *nct* mutants. A similarly targeted transgene with GFP instead of Fz2-C was not able to suppress the defects (Fig. 5C, E, H, J, L-P), demonstrating the specificity of Fz2-C. This full suppression of the phenotypes by nuclear Fz2-C import (Fig. 5A’’) demonstrates that the major relevant event underlying the postsynaptic maturation phenotypes associated with γ-secretase loss is the absence of nuclear Fz2-C.

**Fig. 5.**
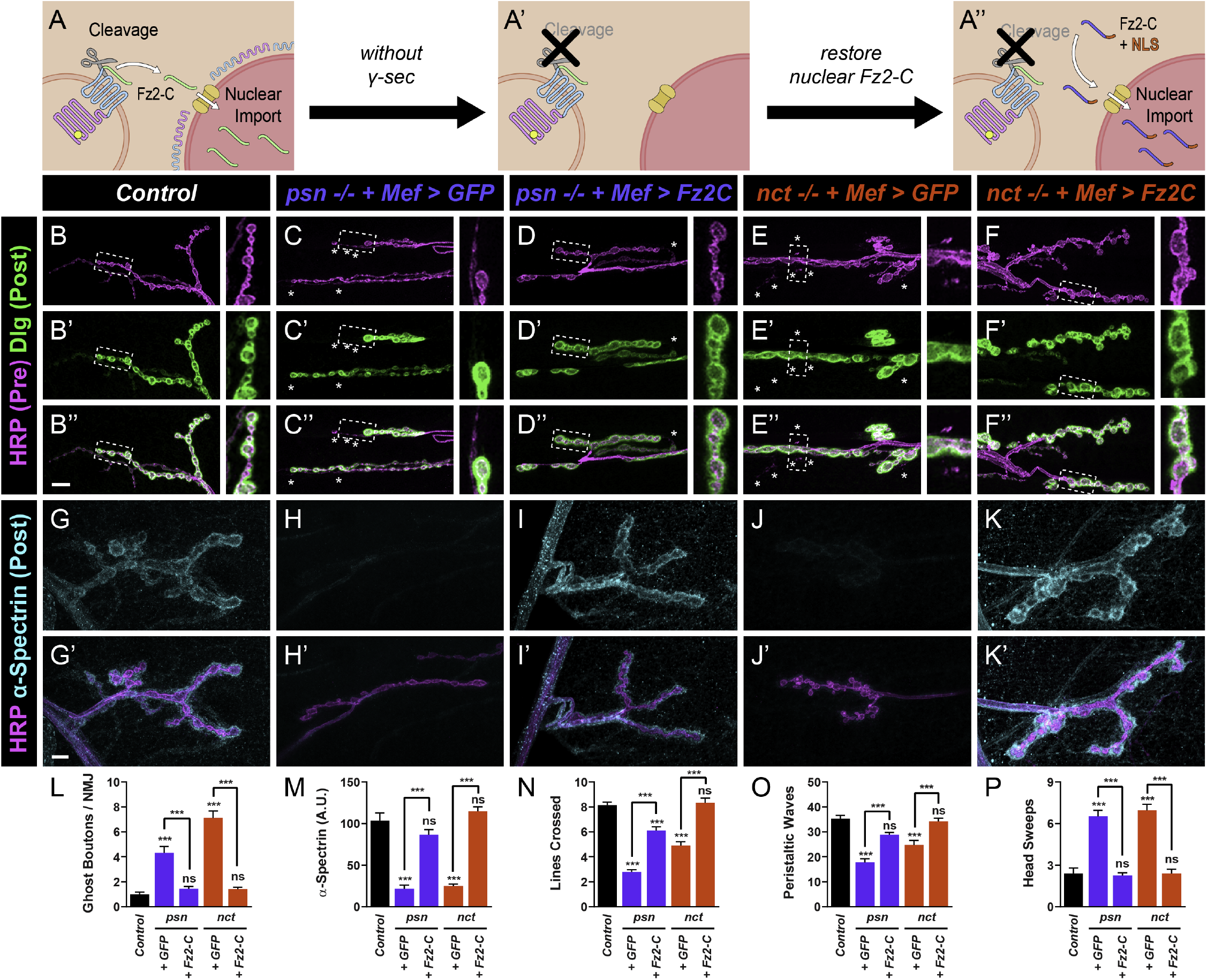
Restoration of nuclear Fz2-C suppresses the morphological and behavioral phenotypes in *psn* and *nct* mutant. (**A**) Diagram of experimental logic. In the normal FNI pathway, Fz2-C cleavage enables nuclear entry, leading to normal postsynaptic development. When cleavage is blocked in γ-secretase mutants, Fz2-C nuclear entry is prevented, resulting in impaired development. If pre-cleaved Fz2-C is expressed in muscles and localizes to the nucleus, it should bypass the need for cleavage, and restore normal development. (**B – F**) Representative confocal images of control larvae (B), *psn* mutant larvae expressing NLS-GFP (C) or NLS-Fz2-C (D) in muscles, and *nct* mutant larvae expressing NLS-GFP (E) or NLS-Fz2-C (F) in muscles and stained with antibodies to HRP (magenta) and Dlg (green). Insets represent high magnification ghost or control boutons marked by the dashed line. Muscle expression of NLS-Fz2-C suppresses the increased incidence of ghost boutons in *psn* and *nct* mutants while NLS-GFP expression has no effect. (**G – K**) Representative confocal images of control larvae (G), *psn* mutant larvae expressing NLS-GFP (H) or NLS-Fz2-C (I) in muscles, and *nct* mutant larvae expressing NLS-GFP (J) or NLS-Fz2-C (K) in muscles and stained with antibodies to HRP (magenta) and α-spectrin (cyan). Muscle NLS-Fz2-C expression also suppresses the defects in α-spectrin fluorescence and thickness. (**L**) Quantification of ghost boutons. (**M**) Quantification of α-spectrin fluorescence. Both graphs show phenotypic suppression as above. (**N – P**) Quantification of lines crossed (N), peristaltic waves (O), and head sweeps (P) in the genotypes above, indicating that NLS-Fz2-C also suppresses *psn* and *nct* mutant behavioral phenotypes. **, *p* < 0.01, ***, *p* < 0.001, n.s. = not significant. For morphology, *n* ≥ 8 larvae, 16 NMJs. For behavior, *n* ≥ 36 larvae per genotype. Scale bar = 10 μm.

### γ-secretase Perturbation affects Central Synapse Development and Maturation

With our discovery that γ-secretase promotes synaptic development and maturation at peripheral NMJ synapses (Fig. 3, S3), we next wondered if that role was conserved at central synapses and evolutionarily across species. As loss of γ-secretase components are lethal prior to adulthood in *Drosophila*, we used RNAi against individual subunits of γ-secretase, expressed specifically in olfactory receptor neurons (ORNs) and quantified active-zone number in the *Drosophila* antennal lobe (*57*). In the antennal lobe, mature synaptic organization is marked by the establishment of a stable active zone number by 10 days post eclosion (*57*). We quantified active zone number in ORNs using Brp-Short-mStraw (Fig. S8A-D) (*57, 58*) and neurite volume using mCD8-GFP (Fig. S5E-H) followed by 3D rendering (*57*) along with RNAi against *psn, nct*, or *aph-1*. In both males and females, there was a 17-23% reduction in the number of Brp-Short puncta (Fig. S8I, K) but normal neurite volume (Fig. S8J, L). This data suggests that at fly central synapses, γ-secretase is also required to establish the mature active zone number, but not for general neuronal growth, consistent with analogous processes at the NMJ.

In mammalian systems, γ-secretase promote mammalian dendritic spine development via EphA4 and Neuregulin (*41, 59*) as well as neuroprotection via ephrinB / EphB2 and TrkB internalization (*60*) and cell-cell contacts via E-cadherin (*61, 62*). As it can localize to cell-cell adhesion points like synapses and potentially play a role in development, we sought to determine if it would similarly promote dendritic spine morphogenesis from filopodia to mushroom-headed spines, a process commonly associated with maturation (*63*). To investigate this possibility, we utilized pharmacological blockade of y-secretase activity using L645,458, which impairs maturation at the fly NMJ (Fig. S5), on primary neuronal cultures. We cultured mammalian cortical neurons in the presence or absence of L645,458 and assayed dendritic spine density and sub-type after 21 days in vitro (DIV). L645,458 application modestly reduced dendritic spine density (Fig. S9A-E) but significantly altered the types of dendritic protuberances observed. In both samples, we observed three major morphological classes: stubby spines, thin filopodia, and mushroom-headed spines. In L645,458-treated samples, there were significantly fewer mature mushroom-headed spines and a concomitant increase in thin filopodia (Fig. S6G). These data suggest that perturbing γ-secretase activity also impairs dendritic and synaptic development at central synapses in both vertebrates and invertebrates.

### Postsynaptic Maturation Defects are Shared by Diverse Neurodegenerative Disease Models

Our discovery of a new role for *presenilin* in promoting synaptic maturation motivated a translational question: given the genetic links between *presenilin* and familial Alzheimer’s Disease (*32, 33*) and the newly discovered role of *presenilin* in promoting maturation, could human *PSEN1* mutations associated with Alzheimer’s impact maturation? Further, do other neurodegenerative disease models display similar maturation defects, suggesting a developmental component to classical models of neurodegenerative disease and providing mechanistic insight into their progression and potential treatment. Current *Drosophila* Alzheimer’s models use overexpression of pathogenic forms of Tau or Aβ consistent with accumulations in humans (*64*) or neuron-specific RNAi (*65*). However, these approaches do not address how patient mutations in *presenilin* influence disease progression, or if loss-of-function recapitulates the phenotypes of Alzheimer’s-linked mutations (*35, 66*). It is further unclear if pathogenic accumulation is causative in Alzheimer’s given recent clinical trial failures (*37*).

We first sought to examine these questions by using CRISPR/Cas9 genome editing to make *Drosophila* lines with *presenilin* mutations equivalent to known human mutations associated with early-onset Alzheimer’s Diseases (EOAD). Such a strategy introduces precise changes into the genome and allows us to examine how disease-associated point mutations expressed under the control of their endogenous locus affect synaptic maturation. We compared human *PSEN1* and *Drosophila psn* and found that of 36 mutations found in EOAD families (*36*), 27 of those were in residues conserved in *Drosophila* Psn. Specifically, a G206D mutation (*36*) with definite pathology and particularly early age of onset corresponded to G228 in *Drosophila psn*. Using CRISPR/ Cas9 (*67*), we introduced the G228D mutation into *Drosophila*

*presenilin* and examined synaptic maturation. *psn^G228D^* mutants were homozygous viable and the larvae displayed a 5-fold increase in ghost boutons (Fig. 6A-B, I), indicating impaired maturation. Such a phenotype is consistent with perturbing other AD susceptibility genes in the fly (*68*). Thus, *presenilin* mutations associated with EOAD in humans also cause failures in postsynaptic maturation and development.

**Fig. 6.**
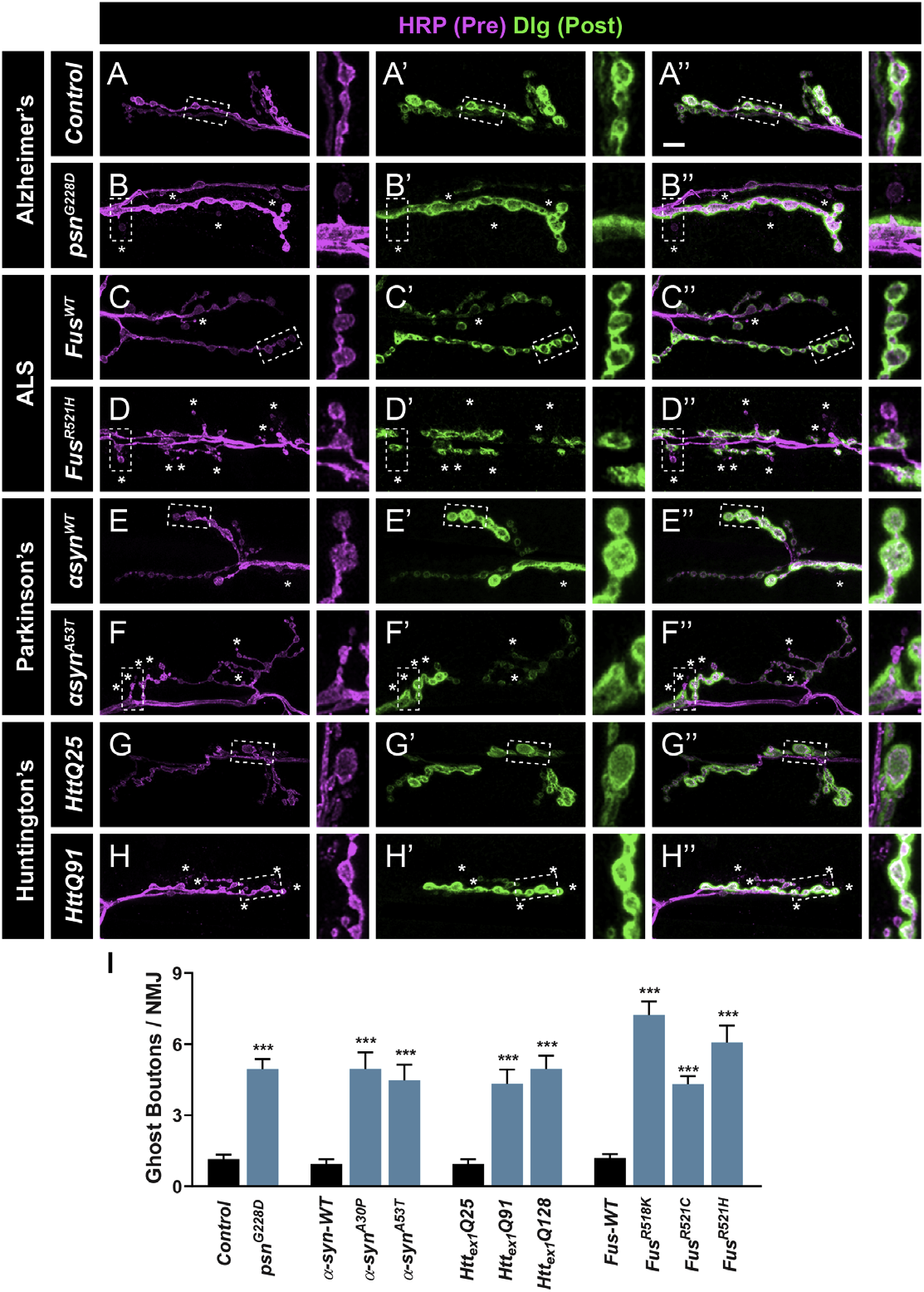
Diverse neurodegenerative disease models display defects in postsynaptic development and maturation. (**A - H**) Representative confocal images of various genotypes stained with antibodies to HRP (magenta) and Dlg (green). Insets represent high magnification ghost or control boutons marked by the dashed line. (**A – B**) Larvae bearing analogous Alzheimer’s disease patient-associated mutations in PSEN1 show an increase in ghost boutons over control larvae. (**C – D**) Neuronal expression of the R521H mutant variation of human Fus from amyotrophic lateral sclerosis patients shows an increase in ghost boutons over expression of wild-type human Fus. (**E – F**) Neuronal overexpression of Parkinson’s disease-associated mutations in human α-synuclein also increase ghost bouton occurrence while wild-type α-synuclein expression does not. (**G – H**) Neuronal expression of pathogenic human Huntingtin (Htt)-repeat containing constructs also result in more ghost boutons while expression of non-pathogenic Htt has normal postsynaptic development and maturation. (**I**) Quantification of ghost boutons in control and neurodegenerative disease model genotypes. For all experiments, *n* ≥ 8 larvae, 16 NMJs. ***, *p* < 0.001. Scale bar = 10 μm.

We next wondered if these defects were specific to Alzheimer’s Disease models, or if synaptic maturation errors were shared among multiple neurodegenerative diseases. We examined 3 additional neurodegenerative disease models in *Drosophila* encompassing amyotrophic lateral sclerosis (ALS), Parkinson’s Disease (PD), and Huntington’s Disease (HD); these models have been exceedingly informative for using *Drosophila* to understand cellular mechanisms of neurodegeneration. (*69*). In each condition, we employed pan-neuronal overexpression of a wild-type human transgene as the control and overexpression of pathogenic forms as the experiment. Intriguingly, each model displayed a significant increase in ghost boutons, suggesting impaired postsynaptic maturation. Overexpression of pathogenic Fus (*70*) from ALS models, α-synuclein from PD models (*71*), and Htt from HD models (*72, 73*) all induced a 5- to 8-fold increase in ghost boutons (Fig. 6C-I). Importantly, overexpression of a wild-type version of each transgene did not impair maturation (Fig. 6C-I), indicating that only the disease-causing pathogenic form perturbed development. This data suggests that, even though these diverse neurodegenerative disease models have different genetic underpinnings, they share similar early developmental phenotypes in perturbed postsynaptic development. As such, understanding the mechanisms that underlie synaptic maturation may offer fundamental insight into our understanding of neurodegeneration.

## Discussion

Synaptic maturation is an essential developmental process that enables the transition from a nascent, unreliable synapse to a robust connection capable of high-fidelity neurotransmission. Here we find an unexpected neurodevelopmental role for γ-secretase, a well-known protein complex closely linked to Alzheimer’s Disease, in promoting postsynaptic development and maturation. γ-secretase functions postsynaptically to enable structural maturation, proper synaptic and cytoskeletal morphology, and coordinated behavior. The γ-secretase complex functions by enabling the cleavage of a Wnt receptor, Fz2, so that the C-terminus translocates to the nucleus, promoting postsynaptic development. Finally, and surprisingly, we find that genetic models of neurodegenerative diseases, including ALS, Alzheimer’s, Parkinson’s, and Huntington’s, display similar neurodevelopmental defects in synaptic maturation.

Our identification of γ-secretase as required for Fz2 cleavage in synaptic development answers a long-standing question about the protease responsible and offers evidence that the Frizzled2 nuclear import pathway primarily promotes postsynaptic development rather than synaptic growth (*23*). This is a critical step forward in our understanding of the molecular mechanisms that underlie synaptic maturation. As maturation defects are associated with neurodevelopmental disorders (*9*), treating these conditions requires a knowledge of the processes that promote the correct maturation of synapses. By first identifying protein complexes, like γ-secretase, that are required for these fundamental neuronal events, we can subsequently examine the regulatory events, cellular partners with which it interacts, and other proteases it may synergize with, to promote maturation. As γ-secretase promotes central and peripheral synapse development in vertebrates and invertebrates (this study) (*42, 43*), it allows us to study how a core developmental requirement for γ-secretase is conserved evolutionarily despite the different roles of each synapse.

More broadly, identifying a novel neurodevelopmental role for γ-secretase and its mechanistic basis offers unique insight into the potential function of γ-secretase as a disease-linked complex. Mutations in PSEN1, the catalytic subunit of γ-secretase, are strongly linked to early-onset Alzheimer’s Disease (*32*) but we lack thorough molecular understanding of this genetic link. With a firmer grasp of the normal role of γ-secretase in basic neuronal biology, we can begin to connect it to the cellular processes that influence neurodegeneration, development, and neuronal function. Such a mechanistic discovery can potentially provide earlier hallmarks to assess disease progression and inform therapeutic strategies aimed at perturbing specific biological pathways (as in Fig. 5) to suppress extant phenotypes or reduce disease severity.

We also observed comparable synaptic defects in multiple neurodegenerative disease (NDD) models including ALS, Parkinson’s, and Huntington’s. This suggests these diseases, with diverse mechanisms, etiologies, progressions, and symptoms, may share a previously unappreciated developmental component. One of the prevailing mysteries surrounding neurodegenerative diseases involves its onset. Though patients carry gene mutations all their lives, why do NDDs manifest when they do, often later in life? The presence of defects associated with synaptic maturation presents a tantalizing hypothesis: if these disease genes are involved in synaptic maturation, the first reflection of mutations in those genes may result in immature synapses that are not properly constructed. They are still capable of function but may lack the longevity of synapses where maturation is normal. As such, in advanced age, synapses that were not constructed correctly are the first synapses to fail, leading to neurodegeneration. Intriguingly, some disease models of Alzheimer’s and Parkinson’s show specific reductions in postsynaptic protein levels that precede neurodegeneration (*74, 75*). As growing evidence suggests that neurodegenerative diseases have synaptopathic origins (*76, 77*), it is intriguing that other synaptic molecules, like Ephs/Ephrins, are modified by γ-secretase (*78*), potentially underlying additional neurodegenerative mechanisms. Synaptic maturation may provide a unique angle to consider neurodegeneration, leading to biomarkers and hallmarks detectable earlier than current clinical approaches. With γ-secretase, maturation defects can be suppressed by activating the downstream pathway (Fig. 5), further suggesting that these approaches may have clinical relevance. Understanding how synaptic maturation influences NDD onset or severity will inform clinical strategies to combat these devastating diseases and allow a better understanding of the basic mechanisms underlying nervous system formation.

## Acknowledgments

We’d like to thank the frontline workers, grocery store employees, drivers, and delivery people who selflessly risked their safety to keep society safer during the COVID-19 pandemic. We also thank Shernaz Bamji, Matthew Dalva, Kristen Davis, Chris Doe, Jesse Humenik, Irwin Levitan, Craig Montell, Joaquin Navajas Acedo, and S. Zosimus for insightful comments on the manuscript. We deeply appreciate the generous gifts of reagents from Jeffrey Axelrod, Hugo Bellen, Vivian Budnik, Matthew Dalva, Graeme Davis, Mark Fortini, Alison Goate, Irwin Levitan, Le Ma, Kate O’Connor-Giles, Margaret Pearce, and Thomas Schwarz. Stocks obtained from the Bloomington Drosophila Stock Center (NIH P40OD018537) were also used in this study. TJM would especially like to thank Thomas Schwarz, in whose lab his fascination with Fz2-C began. This work was supported by US National Institute of Health grants R01-NS110907 and R00-DC013059 (to TJM). Work in the TJM Lab is supported by grants from the Alfred P. Sloan Foundation, the Whitehall Foundation, the Jefferson Dean’s Transformational Science Award, the Jefferson Synaptic Biology Center, the Commonwealth Universal Research Enhancement program of the Pennsylvania Department of Health, and Thomas Jefferson University start-up funds.

## Author Contributions

L.R. and T.J.M. designed the project; L.R., A.D., E.M., M.A., J.C.D., S.T., H.F., and T.J.M. performed experiments; M.P. and T.J.M. produced reagents; L.R., A.D., E.M., M.A., and T.J.M. analyzed the data; L.R. and T.J.M. wrote the manuscript; L.R., A.D., E.M., M.A., J.D., M.P., and T.J.M. edited the manuscript.

## Materials and Methods

### RNAi protease screen

Fz2 cleavage occurs at a consensus (KTLES) glutamyl-endopeptidase site (*23*). We used two GO_MOLECULAR_ FUNCTION terms in FlyBase (*80*): “endopeptidase” and “metalloprotease” and identified 121 total genes with those predicted molecular functions. From that list, 116 genes had 1 or more RNAi lines available at the Vienna Drosophila Resource Center (VDRC) or the Harvard TRiP Collection (*26, 81*). We omitted the Neprilysin family (26 genes) as its active site faces the extracellular space (*82*) and DFz2-C cleavage is expected to occur in the cytoplasm. We obtained RNAi lines for the remaining 90 candidates as well as 3 controls: two positive controls - dfz2 (*23*) and *trol* (*20*) – known to impair postsynaptic development and maturation, and one negative control – GFP. Because Fz2 cleavage occurs in the muscle, we conducted the screen by driving each RNAi line specifically in muscles using the DMef2-GAL4 driver (*83*). We reasoned that impairing Fz2 cleavage (by blocking the relevant protease) would impair postsynaptic development and maturation similarly to the removal of the Fz2 receptor itself and show similar phenotypes. We screened F1 progeny of crosses between each RNAi line and the DMef2-GAL4 driver (with the experimenter blind to genotype) and quantified “ghost boutons,” a hallmark of impaired postsynaptic maturation (*13, 14, 16, 21, 53, 84, 85*). Positive hits were lines that caused a significant increase in ghost boutons over the negative control. For all genotypes, we performed immunocytochemistry in third instar larvae (see below) and stained with antibodies against postsynaptic Dlg and presynaptic HRP. For all genotypes, at least 6 NMJs in 6 larvae were scored. To ensure that any potential defects were not due to destabilization of the synapse, we also scored “footprints,” which are retracted synapses that are positive for Dlg staining and negative for HRP staining (*27–29*). Only four genes showed a significant increase in ghost boutons over the negative control: *dfz2* and *trol* (the positive controls), *psn*, and *nct*.

### Drosophila stocks and transgenic strains

All control genotypes, mutant combinations, transgenic lines, manipulations, and crosses were maintained on cornmeal medium (Archon Scientific, Durham, NC) at 25°C and 60% relative humidity with a 12/12 light/dark cycle. *N^TS^* mutants were raised as described (*86*). All genetic components (mutant alleles, transgenes, drivers, etc.) were maintained over larvally selectable balancer chromosomes to enable facile identification at the appropriate life stage. The following mutant alleles were used: *psn^143^* (*87*), *psn^C4^, psn^I2^* (*88, 89*), *nct^A7^, nct^E3^, nct^J2^* (*90*), *aph-1^D35^* (*91*), *Df(2L)Exel6277* (as *aph-1^Df^*) (*92*), *pen-2^MI02639^* (*93*), *dfz2^C1^* (*94*), Df(3L)ED4782 (as *dfz2^Df^*) (*23*), *appl^D^* (*95*), *N^ts1^* (*86, 96*), *msk^5^* (*97*), *imp-β11^70^* (*98*), Df(2R)Δm22 (as *imp-β11^Df^*) (*98, 99*), *Df(X)ten-a* (*53*). The following UAS transgenes were used: *UAS-Fz2-FLAG* (*56*), *UAS-Psn-Nmyc, UAS-Nct-2myc, UAS-Aph1-V5, UAS-Pen2-2flag* (*45*), *UAS-Fz2-FL, UAS-Fz2-ΔKTLES, UAS-myc-NLS-Fz2-C* (*23*), *UAS-NLS-GFP* (*100*), *UAS-DshDIX* (*25*), *UAS-kuz-DN* (*101*), *UAS-psn-IR-43082, UAS-aph-1-IR-16820, UAS-mam-IR-102091* (*26*), *UAS-nct-IR-JF02648, UAS-pen-2-IR-JF02608* (*81*), *UAS-Fus WT, UAS-Fus R518K, UAS-Fus R521C, UAS-Fus R521H* (*70*), *UAS-WT α-synuclein, UAS-A30P α-synuclein, UAS-A53T α-synuclein* (*71*), *UAS-HttQ25-YFP, UAS-HttQ91-YFP* (*72*), *UAS-Htt-Q103* (*73*). The following GAL4 driver lines were used to achieve tissue-specific expression: *elavC^155^-GAL4* (*102*) or *elav-GAL4* (*103*) (pan-neuronal expression), *OK371-GAL4* (*104*) (motoneuron-specific expression), *DMef2-GAL4* (*83*) (pan-muscle expression) *Or67d-GAL4* (*105*) (DA1 ORN expression), *Or47b-GAL4* (*106*) (VA1lv ORN expression), *pebbled-GAL4* (*107*) (pan-ORN expression).

### Construction of psn^G228D^

The sequences of human Presenilin1 (PSEN1) and *Drosophila* Presenilin (Psn) were aligned using SnapGene software (Insightful Science, San Diego, CA) and the equivalent residues from known patient mutations (*36*) identified. Psn G228D was selected as the *Drosophila* equivalent to human PSEN1 G206D. We used CRISPR/Cas9 editing (*108*) with WellGenetics, Inc. (Taipei City, Taiwan) to make a custom-designed gRNA and accompanying construct to introduce the specific mutation. Four lines were obtained and sequenced to confirm the presence of the change. One line bearing the mutation was stabilized over a third chromosome balancer and used for further experiments.

### Cloning of HA-Fz2-FLAG constructs and transgenic line production

The HA- and FLAG-tagged Fz2 construct has a 3X HA tag inserted at amino acid position 220 (between the cysteine rich domain and the transmembrane domain) and a C-terminal 3X-FLAG tag. Previous work suggested those locations did not interfere with normal protein function (*56, 109*). These were inserted into sequence amplified from a full length BDGP-Gold Fz2 cDNA clone (LD10629 in pBluescript) from the *Drosophila* Genomics Resource Center (DGRC, supported by NIH grant 2P40OD010949). All PCR amplification steps used the high fidelity Q5 Polymerase (New England Biolabs) and primers from Integrated DNA Technology (Coralville, IA). The FLAG tag was amplified from pUAST-W-tFHAH-attB (*53*) and cloned into LD10629 which was linearized and assembled using In-Fusion HD (Takara Bio). This resulted in plasmid pBS-FZ2.FLAG which was used to amplify the dFZ2 ORF with the FLAG tag for cloning into the Gateway entry vector pENTR-D-TOPO (Thermo Fisher). This step eliminated most of the dFZ2 5’ and 3’ UTRs while maintaining the presumptive Kozak sequence adjacent to the start codon. Following In-Fusion assembly, the resulting plasmid, pENTR-FLAG#5.ORF, was used to engineer the HA tag between the dFZ2 cysteine rich and transmembrane domains. The HA epitope tag (amplified from pUAST-W-tFHAH-attB) was inserted by In-Fusion cloning of multiple fragments in pENTR-FLAG#5.ORF. Positive clones were screened and verified by PCR diagnostics and Sanger sequencing. This gave pENTR-HA-FZ2-FLAG which was then used in a Gateway LR reaction with the pUAST-Gateway-attB vector to generate pUAST-HA-FZ2-FLAG. This plasmid was used to generate transgenic flies (Bestgene, Chino Hills, CA) with the construct integrated into the VK00037 docking site.

### Cleavage assay, Western blot, SDS-PAGE analyses

UAS-Fz2-FLAG was expressed in muscles using *DMef2-GAL4* in a wild-type, *imp-β11* mutant, *psn* mutant, or *nct* mutant background. Lysates were prepared from partially dissected third instar larval body walls as described (*16*). Proteins were separated on 4-15% Mini-PROTEAN TGX gels (BioRad, Hercules, CA) and transferred to nitrocellulose. Blots were incubated overnight in primary antibodies at 4°C and secondary antibodies at room temperature (21-22°C) for 1 hour. The following primary antibodies were used: mouse anti-FLAG M2 (Sigma-Aldrich, cat. no. F1804, 1:5000) and mouse anti-α-tubulin DM1a (Sigma-Aldrich, cat. no. T9026, 1:10000). HRP-conjugated secondary antibodies were used at 1:10000 (Jackson ImmunoResearch, West Grove, PA). Blots were developed using the SuperSignal West Femto Maximum Sensitivity Substrate Kit (ThermoFisher Scientific, Waltham, MA).

### Production of Nicastrin antibodies

Custom antibodies were made against a TSKDFTQLTEVNDFKSLNPDSLQ-C peptide corresponding to amino acids 462-484 of the Nicastrin protein (Pacific Immunology Corp., Ramona, CA). Rabbit antisera were affinity-purified against the original immunizing peptide and used at a dilution of 1:500 on wandering third instar larvae (see below). Specificity of the antisera was validated by the absence of signal in *nct^A7/J2^* mutant larvae.

### Immunocytochemistry

Wandering third instar larvae were dissected and stained as described (*16*). The following primary antibodies were used: mouse anti-Dlg (DSHB, cat. no. mAb4F3, 1:500) (*110*), rabbit anti-Presenilin (custom, 1:200) (*111*), rabbit anti-Nicastrin (custom, 1:500) (this study), rabbit anti-Dlg (custom, 1:40000) (*112*), mouse anti-myc (DSHB, cat. no mAb9E10, 1:100), rabbit anti-myc (ThermoFisher Scientific, cat. no PA5-85185, 1:200), mouse anti-α-spectrin (DSHB, cat. no. mAb3A9, 1:50) (*113*), mouse anti-Brp (DSHB, cat. no. mAbnc82, 1:250) (*114*), rabbit anti-GluRIIC (custom, 1:2500) (*115*), rabbit anti-Syt I (custom, 1:4000) (*116*), mouse anti-CSP (DSHB, cat. no. mAb6D6, 1:100) (*117*), rabbit anti-dsRed (TaKaRa Bio, cat. no. 632496, 1:250 (*57*), chicken anti-GFP (Aves, cat. no. GFP-1020, 1:1000) (*57*), rat anti-N-Cadherin (DSHB, cat. no. mAbDNEX-8, 1:40) (*118*), rabbit anti Fz2-N (custom, 1:100) (*119*), rabbit anti Fz2-C (custom, 1:200) (*119*), mouse anti Lamin C (DSHB, cat. no. mAbLC28.26, 1:200) (*120*), mouse anti-FLAG M2 (Sigma-Aldrich, cat. no. F1804, 1:500), rabbit anti-FLAG (Sigma-Aldrich, cat. no. F7425, 1:250), rabbit anti-Wingless (custom, 1:200) (*55*). Where noted, monoclonal antibodies were obtained from the Developmental Studies Hybridoma Bank, created by the NICHD of the NIH and maintained at The University of Iowa, Department of Biology. Alexa488-, Alexa647- (Jackson ImmunoResearch, West Grove, PA), and Alexa546-conjugated (ThermoFisher, Waltham, MA) secondary antibodies were used at 1:250. FITC-, Cy3-, or Alexa647-conjugated goat anti-HRP primary antibodies were used at 1:100 (Jackson ImmunoResearch, West Grove, PA). Texas-Red-conjugated phalloidin was used at 1:300 (Sigma Aldrich, St. Louis, MO).

### Proximity ligation assay

Larvae were processed as described (*121*) using the DuoLink Mouse Rabbit in situ PLA Kit (Sigma-Aldrich, cat. no. DUO92101, St. Louis, MO). Controls were performed with one or the other epitope-tagged transgene absent or with the probes replaced by water during the first PLA step to ensure that signal observed as not background or bleed-through of channels. Larvae were then imaged via confocal microscopy as described (see below).

### Imaging, parameter quantification, image processing

RNAi screen imaging was conducted on a Leica SP8 confocal microscope (Leica Microsystems, Wetzlar, Germany). All other larvae were imaged with a Zeiss LSM880 with Fast AiryScan confocal microscope (Carl Zeiss, Oberlochen, Germany) using a 10X 0.4 NA, 40X 1.4 NA PlanApo, or a 63X 1.4 NA PlanApo lens. Images were processed and quantified as described (*16, 122*). Ghost boutons were quantified as HRP-positive and Dlg-negative and samples blinded during imaging and quantification. Comparison of fluorescence levels for various parameters was done on samples that were imaged using identical confocal settings, laser powers, and conditions. Images were processed and figures constructed using ZEN 2.3 software (Carl Zeiss, Oberlochen, Germany), Adobe Photoshop 2020, and Adobe Illustrator 2020 (Adobe Systems, San Jose, CA). Fluorescence intensity was measured with ImageJ (NIH, Bethesda, MD).

### Behavioral larval crawling assays

Crawling assays were conducted, and peristaltic waves and head sweeps determined, as described (*123–125*). The experimenter was blind to larval genotype during quantification.

### Pharmacological inhibition of γ-secretase in vivo

Larvae were raised on small-batch *Drosophila* cornmeal dextrose medium (*126*) prepared with 5 μM L685,458 (Tocris Bioscience, Minneapolis, MN) and processed as above. This drug has been used *in vivo* as a potent and specific γ-secretase activity blocker (*43*).

### Primary neuron culture

Dissociated cortical neurons were prepared from embryonic day 17 Long Evans rats (Charles River). Cortices were incubated with 10 μg/mL papain (Worthington Biochemical Corporation) in HBSS for 4 min at 37 °C. Following three washes in HBSS with 0.01 g/mL trypsin inhibitor (Sigma), cortices were triturated with a fire-polished glass Pasteur pipette 5–10 times to obtain a homogeneous cell suspension. Neurons were plated on poly-D-lysine (BD Biosciences, Bedford, MA) and laminin (BD Biosciences)-coated glass coverslips (12 mm; Bellco Glass, Vineland, NJ) in 24-well plates (Corning Life Sciences, Lowell, MA). Neurons were plated at a density of 6 ×10^5^ cells/cm^2^ and cultured in Neurobasal media (Invitrogen, Carlsbad, CA) supplemented with B-27 (Invitrogen), 1% glutamine (Invitrogen), and 1% penicillin–streptomycin (Invitrogen) and maintained in a humidified incubator with 5% CO_2_ at 37°C.

### Neuronal transfection and drug treatment

Neurons were transfected using Lipofectamine 2000 (Invitrogen) after 3 days in vitro. Transfection mixture was prepared (per coverslip) as follows: 0.5 μL Lipofectamine 2000 was added to 50 μL neurobasal medium (without supplement) in a polystyrene tube (USA Scientific). DNA was added to 50 μL neurobasal medium (without supplement) in an Eppendorf tube. After 5 min, the DNA mixture was added to Lipofectamine and incubated at room temperature for 15 min. Conditioned media was removed from the neuronal cultures and replaced with 300 μL of warm neurobasal medium (without supplement). Transfection mixture was added to the neuronal culture and incubated for 2 h at 37 °C and subsequently replaced with filter-sterilized warm conditioned media. For drug treatment, compound L685,458 (Tocris Bioscience, Minneapolis, MN) dissolved in DMSO was added at a final concentration of 2.5 μM once after 7 days in vitro. For the control condition, the same volume of DMSO was added.

### Primary neuron immunocytochemistry

After 21 days in vitro, neurons were fixed with 4% paraformaldehyde and 2% sucrose for 10 minutes. Neurons were washed three times with PBS. For antibody labeling, neurons were blocked in PBS plus 5% goat serum and 0.1% Triton (PGT) for 1 hr and then incubated with primary antibodies (GFP; Aves lab) diluted in PGT overnight at 4°C After washing, neurons were incubated Alexa 488 secondary antibodies (Jackson ImmunoResearch or Invitrogen) diluted in PGT for 2 hr at room temperature. Finally, neurons were washed three times with PBS and mounted with Mowiol.

### Quantification of synaptic parameters in adult Drosophila

Synaptic puncta (Brp-Short-mStraw) and neurite membrane volume (mCD8-GFP) were imaged, processed, and quantified as described (*57*). All images were obtained as above using a 63X 1.4 NA PlanApo lens and quantified / processed using Imaris Software 9.3.1 (Oxford Instruments, Abingdon, UK) on a custom image processing computer (Digital Storm, Fremont, CA).

### Statistical analysis

Statistical analysis was performed and graphical representations prepared using Prism 8.4.3 (GraphPad Software, Inc., La Jolla, CA). Significance between two samples was determined using a two-tailed Student’s t-test; significance amongst 3 or more samples was determined using one-way ANOVA with a Dunnett post-hoc test to a control sample and a Bonferroni post-hoc test amongst all samples. In each figure, unless otherwise noted, statistical significance is denoted in comparison to control genotypes.

### Genotypes

See Table 1 of complete genotypes by figure panel.

---

**Fig. S1.**
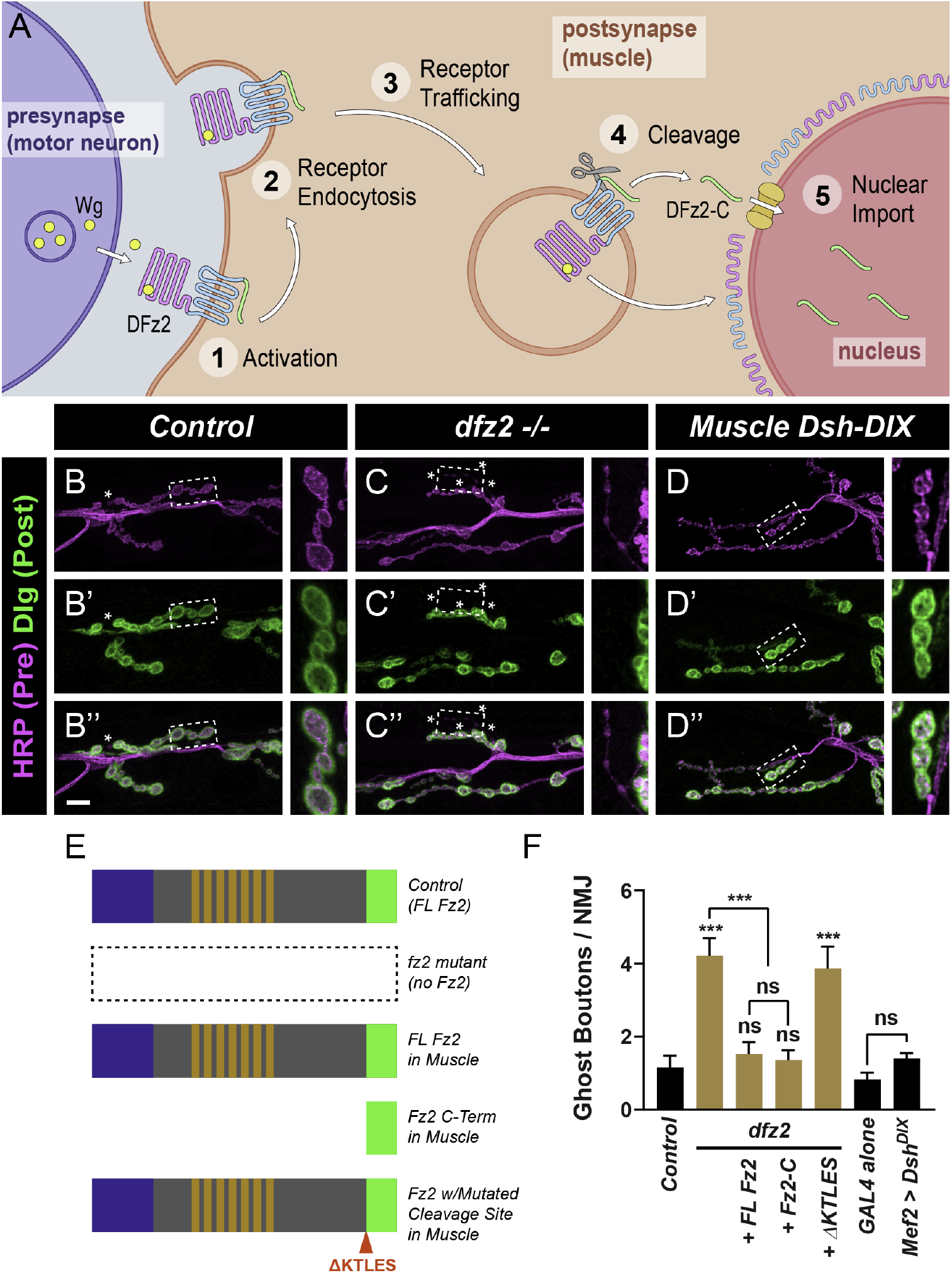
The Frizzled nuclear import pathway promotes postsynaptic development and maturation. (**A**) Schematic of the Frizzled nuclear import pathway (*23*) in *Drosophila*. The Wnt ligand, Wingless, is released from presynaptic motoneurons in an activity-dependent (*13*) fashion where it transverses the synaptic cleft to activate postsynaptic Fz2 receptors. Fz2 is then endocytosed and trafficked to the perinuclear region. There, the C-terminus of the receptor is cleaved and while the N-terminus of the receptor remains localized to the perinuclear space (*23*), Fz2-C is actively imported into the nucleus (*16, 23*) to promote postsynaptic development and maturation. (**B – D**) Representative confocal images of larvae stained with antibodies to HRP (magenta) and Dlg (green). Asterisks indicate ghost boutons. Insets represent high magnification ghost or control boutons marked by the dashed line. *dfz2* mutants (C) show an increased incidence of ghost boutons over control larvae (C) while muscle expressed Dishevelled dominant negative (Dsh-DIX) resembles control (D). This data indicates *fz2* is required for postsynaptic maturation and development but not likely through canonical Dsh signaling. (**E**) Structure-function diagram of the Fz2 constructs used to determine which part of the receptor was necessary for postsynaptic development and maturation. (**F**) Quantification of ghost boutons. Loss of *fz2* increases ghost boutons 4-fold. This phenotype can be suppressed by muscle expression of full-length Fz2 or just the C-terminus, but not by a Fz2 construct where the cleavage site is mutated (*16*). Dsh-DIX expression shows no phenotype. *n* ≥ 6 larvae, 12 NMJs. ***, *p* < 0.001, n.s. = not significant. Scale bar = 10 μm.

**Fig. S2.**
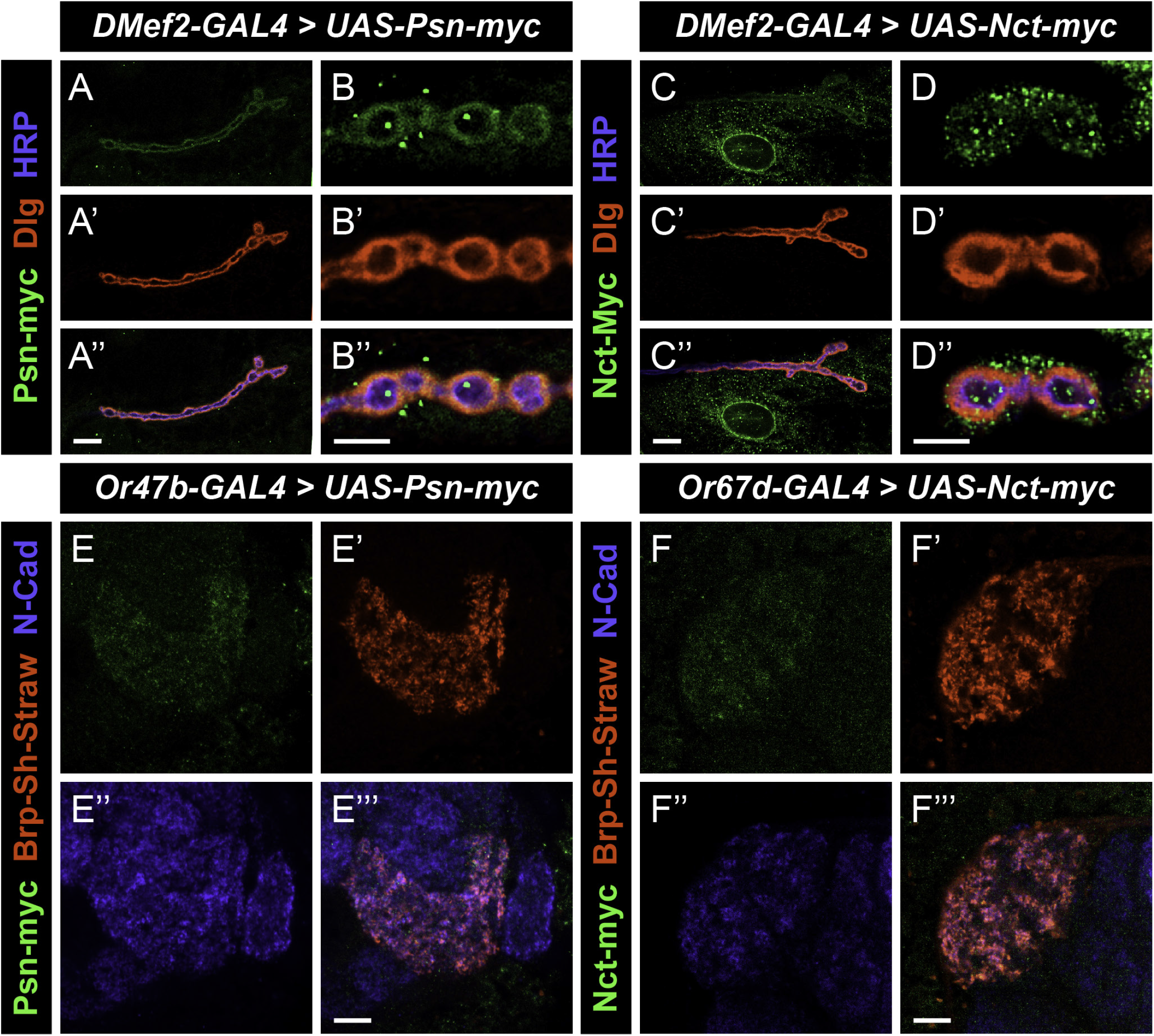
Epitope-tagged Presenilin and Nicastrin can localize to synaptic regions. (**A – D**) Representative confocal images of larvae expressing Psn-myc (A – B) or Nct-myc (C – D) in muscles and stained with antibodies to Myc (green), Dlg (red), and HRP (blue). Exogenously expressed Psn and Nct can localize to the postsynaptic region, consistent with endogenous localization of each protein. High magnification images show an overlap with the largely postsynaptic marker Dlg (B, D). Note also that Nct-myc can also localize to the nuclear periphery (C – D), where Fz2 is cleaved to promote nuclear entry. (**E – F**) Representative single confocal sections of adult brains expressing Psn-myc (E) or Nct-myc (F) and the active zone label Brp-Short in Or47b-positive olfactory receptor neurons that project to the VA1v glomerulus and stained with antibodies to Myc (green), Brp-Short (red), and N-Cadherin (blue). Both Psn-myc and Nct-myc can localize to the synapse-rich regions of these neurons, indicating synaptic localization in both the central and peripheral nervous systems of γ-secretase components. Scale bar = 10 μm (A, C), 5 μm (B, D), or 20 μm (E, F).

**Fig. S3.**
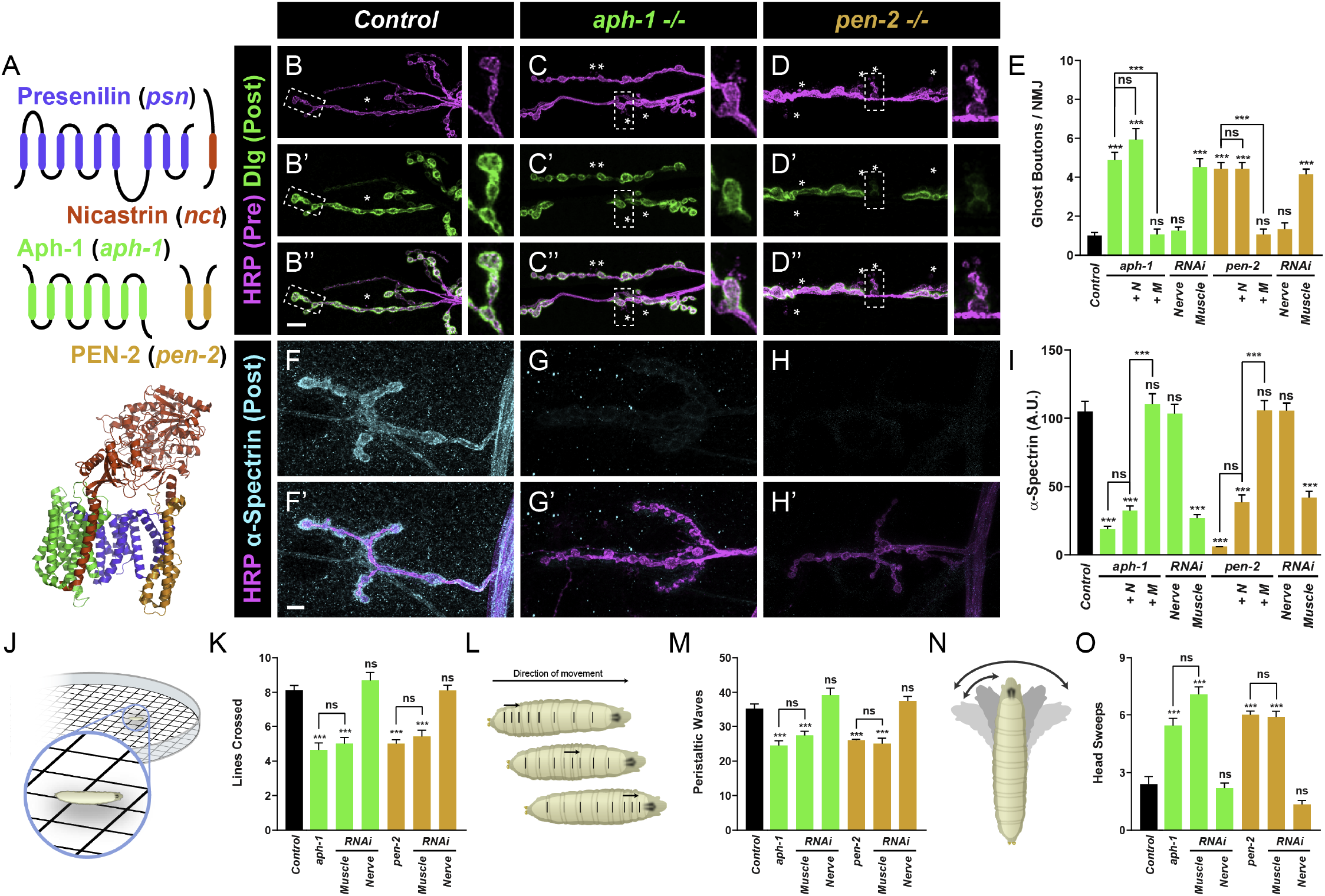
Loss of postsynaptic *aph-1* or *pen-2* impairs postsynaptic development and function. (**A**) Schematic of individual γ-secretase subunits (top): Presenilin (blue), Nicastrin (red), Aph-1 (green), and Pen-2 (orange) and ribbon diagram (bottom) representing the structure of the γ-secretase holocomplex (*79*). (**B – D**) Representative confocal images of control (B), *aph-1* mutant (C), and *pen-2* mutant (D) larvae stained with antibodies against HRP (magenta) and Dlg (green). Asterisks indicate ghost boutons and insets represent high magnification ghost or control boutons marked by the dashed line. *aph-1* and *pen-2* mutants show an increased incidence of ghost boutons, indicating impaired postsynaptic development and maturation. (**E**) Quantification of ghost boutons in mutant, rescue, and RNAi genotypes. *aph-1* and *pen-2* mutants show a 5-fold increase in ghost boutons over control genotypes. The ghost bouton phenotype can be suppressed by postsynaptic muscle expression of an Aph-1 or Pen-2 transgene in the respective mutant (+ M) but not by presynaptic neuronal expression (+ N). Further, the phenotype can be recapitulated by muscle RNAi against *aph-1* or *pen-2* but not neuronal RNAi. (**F – G**) Representative confocal images of control (F), *aph-1* mutant (G), and *pen-2* mutant (H) larvae stained with antibodies against HRP (magenta) and α-spectrin (cyan). α-spectrin fluorescence and thickness is markedly reduced (but not eliminated) in *aph-1* and *pen-2* mutants while HRP staining is unaffected. (**I**) Quantification of α-spectrin fluorescence intensity in mutant, rescue, and RNAi genotypes. α-spectrin is reduced by 80% in *aph-1* and *pen-2* mutants; the α-spectrin phenotype is suppressed by muscle expression (+ M) but not neuronal expression (+ N), indicating a postsynaptic function for each in maturation and development. Similarly, muscle RNAi but not neuronal RNAi induces a comparable phenotype, as with *psn* and *nct*. (**J**) Diagram of larval crawling assay to measure motility. (**K**) Quantification of larval motility in mutant and RNAi genotypes. Loss of *aph-1* and *pen-2* impairs the number of lines crossed. In both mutants, the crawling deficit is completely recapitulated by muscle RNAi and not neuronal, indicating a postsynaptic role in larval motility. (**L**) Diagram of larval peristaltic waves with arrows indicating direction of movement and lines on the larva denoting body wall segments. (**M**) Quantification of peristaltic waves in mutant and RNAi genotypes. Loss of *aph-1* or *pen-2* reduces peristaltic wave number. The peristalsis defect is completely recapitulated in muscle RNAi of either gene but not by neuronal RNAi. (**N**) Diagram of the larval head sweep. Arrows and shading indicate directions of motion during the head sweep behavior. (**O**) Quantification of head sweeps in mutant and RNAi behavior. Both *aph-1* and *pen-2* mutants show an improper 2.5-fold increase in head sweeps over control larvae. This behavioral phenotype is recapitulated again by muscle, and not neuronal RNAi. Taken together, the data suggests that postsynaptic *aph-1* and *pen-2* are essential to promote normal postsynaptic morphology, maturation, and larval behavior. As with *psn* and *nct*, all four subunits of γ-secretase have nearly identical phenotypes when their postsynaptic function is perturbed. For morphological experiments, *n* ≥ 8 larvae, 16 NMJs. For behavioral experiments, *n* ≥ 24 larvae. **, *p* < 0.01, ***, *p* < 0.001, n.s. = not significant. Scale bar = 10 μm.

**Fig. S4.**
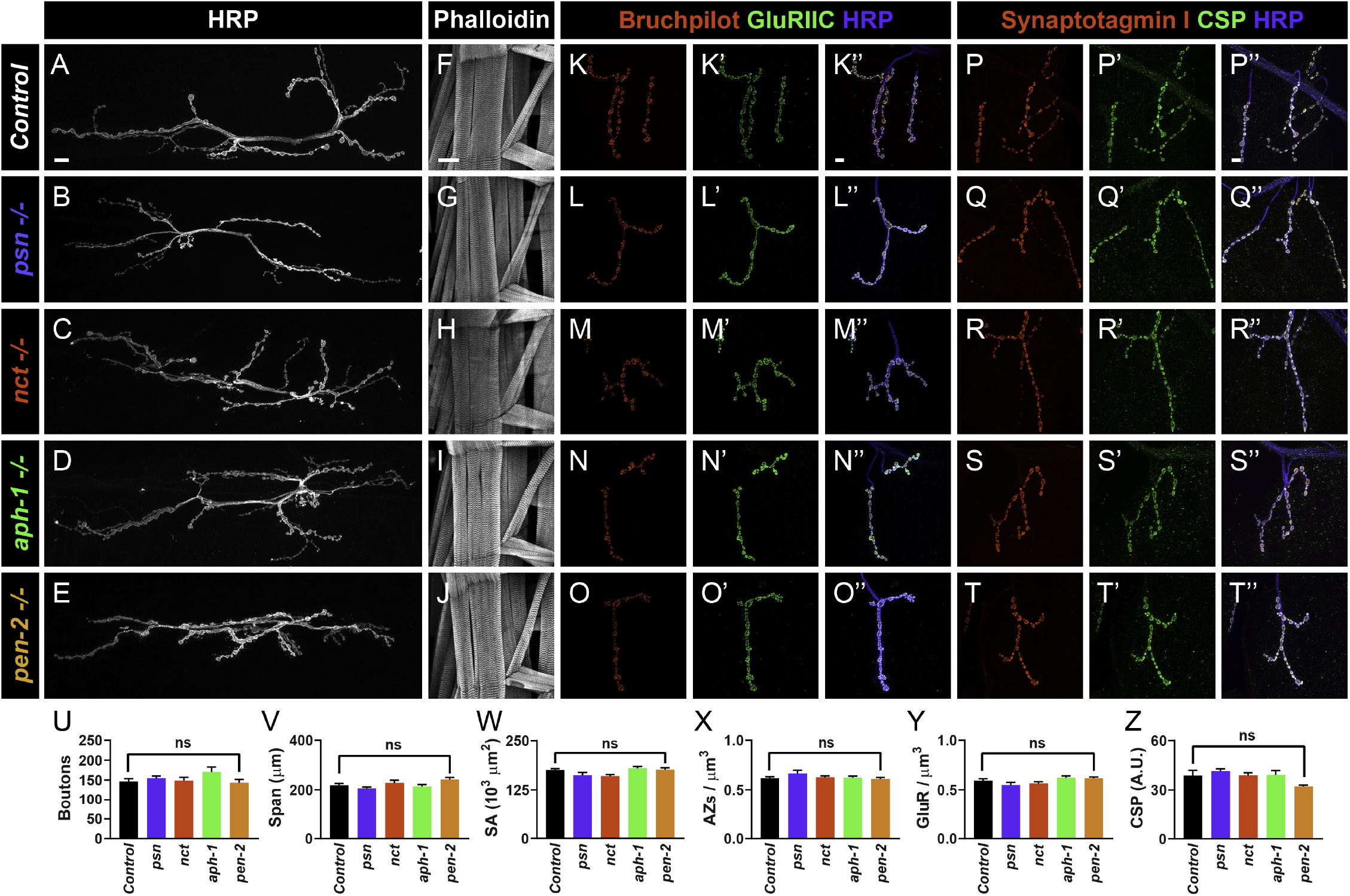
γ-secretase mutants show otherwise normal synaptic development. (**A – E**) Representative NMJs at muscle 6/7 stained with antibodies to HRP in control (A), *psn* mutant (B), *nct* mutant (C), *aph-1* mutant (D), and *pen-2* mutant (E) larvae. (**F – J**) Representative images of the larval body wall muscle field containing muscles 6, 7, 12, 13, 5, 8, and 4 stained with TexasRed-conjugated phalloidin in control (F), *psn* mutant (G), *nct* mutant (H), *aph-1* mutant (I), and *pen-2* mutant (J) larvae. (**K – O**) Representative NMJs at muscle 4 stained with antibodies to Bruchpilot (red), GluRIIC (green), and HRP (blue) in control (K), *psn* mutant (L), *nct* mutant (M), *aph-1* mutant (N), and *pen-2* mutant (O) larvae. (**P – T**) Representative NMJs at muscle 4 stained with antibodies to Synaptotagmin I (red), CSP (green), and HRP (blue) in control (P), *psn* mutant (Q), *nct* mutant (R), *aph-1* mutant (S), and *pen-2* mutant (T) larvae. In all cases, each mutant parameter appears similar to control larvae, suggesting no defects in generalized synaptic development. (**U – Z**) Quantification of bouton number (U), NMJ length / synaptic span (V), muscle surface area (W), active zone density (X), glutamate receptor density (Y), and CSP fluorescence (Z) in all genotypes. No quantitative differences or phenotypes were observed in any of the γ-secretase subunit mutants. This data indicates that defects in postsynaptic development and maturation are unlikely to be secondary to gross defects in synapse formation and organization. In all cases, *n* ≥ 6 larvae, 12 NMJs. n.s. = not significant. SA = surface area. AZs = active zones. GluR = glutamate receptor. Scale bar = 10 μm (A – E, K – T) or 100 μm (F – J).

**Fig. S5.**
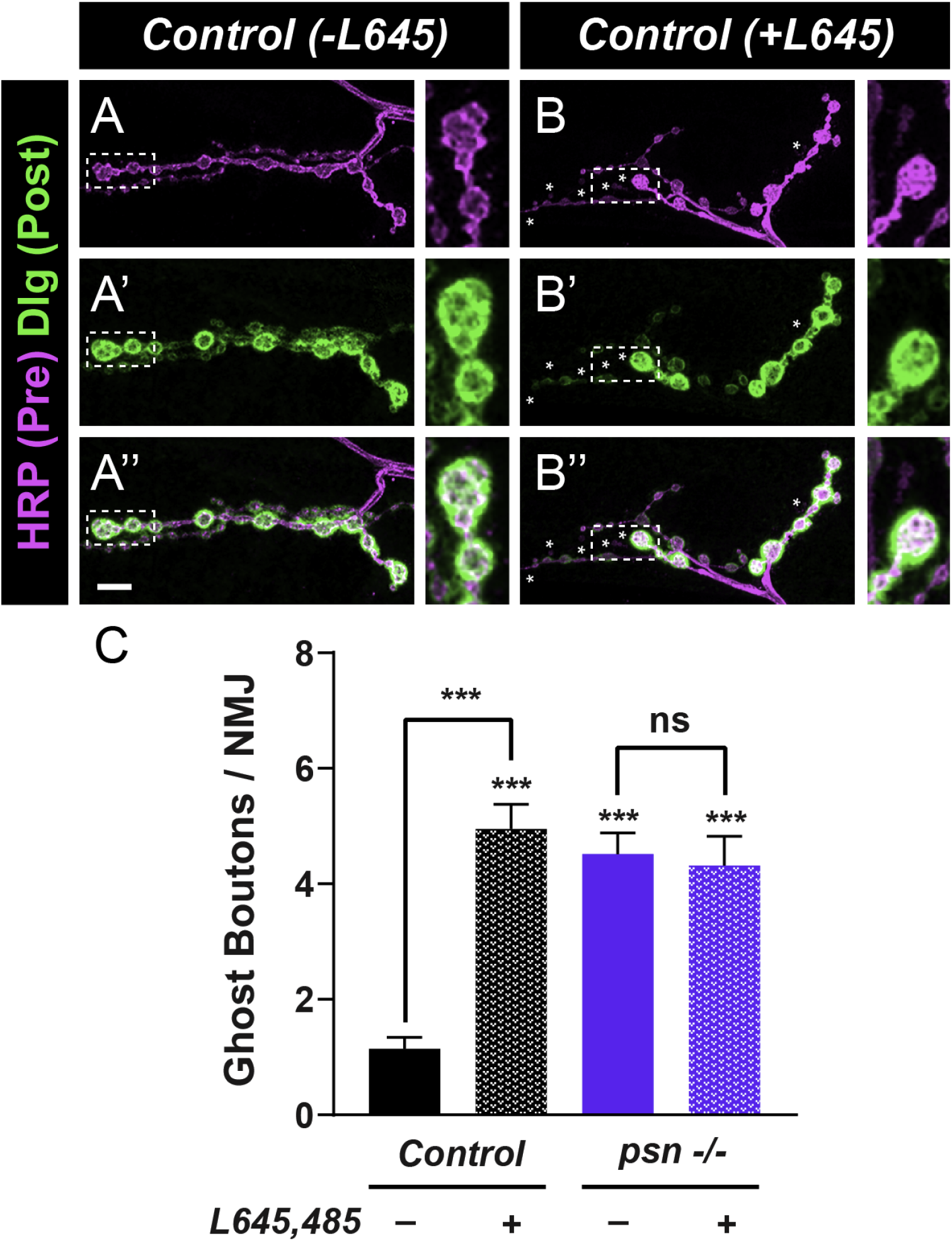
Pharmacological blockade of γ-secretase activity in *Drosophila* impairs postsynaptic development. (**A – B**) Representative confocal images of non-drug treated (A) and drug-treated (B) larvae stained with antibodies for HRP (magenta) and Dlg (green). Asterisks indicate ghost boutons. Insets represent high magnification ghost or control boutons marked by the dashed line. Larvae reared on L645,485 show increased ghost boutons. (**C**) Quantification of ghost boutons. Rearing on L645,485 causes a 5-fold increase in ghost boutons in otherwise-wild-type larvae. This change is not observed in *psn* mutants, suggesting that these do not enhance each other and likely target the same pathway. In all cases, *n* ≥ 8 larvae, 15 NMJs. ***, *p* < 0.001, n.s. = not significant. Scale bar = 10 μm.

**Fig. S6.**
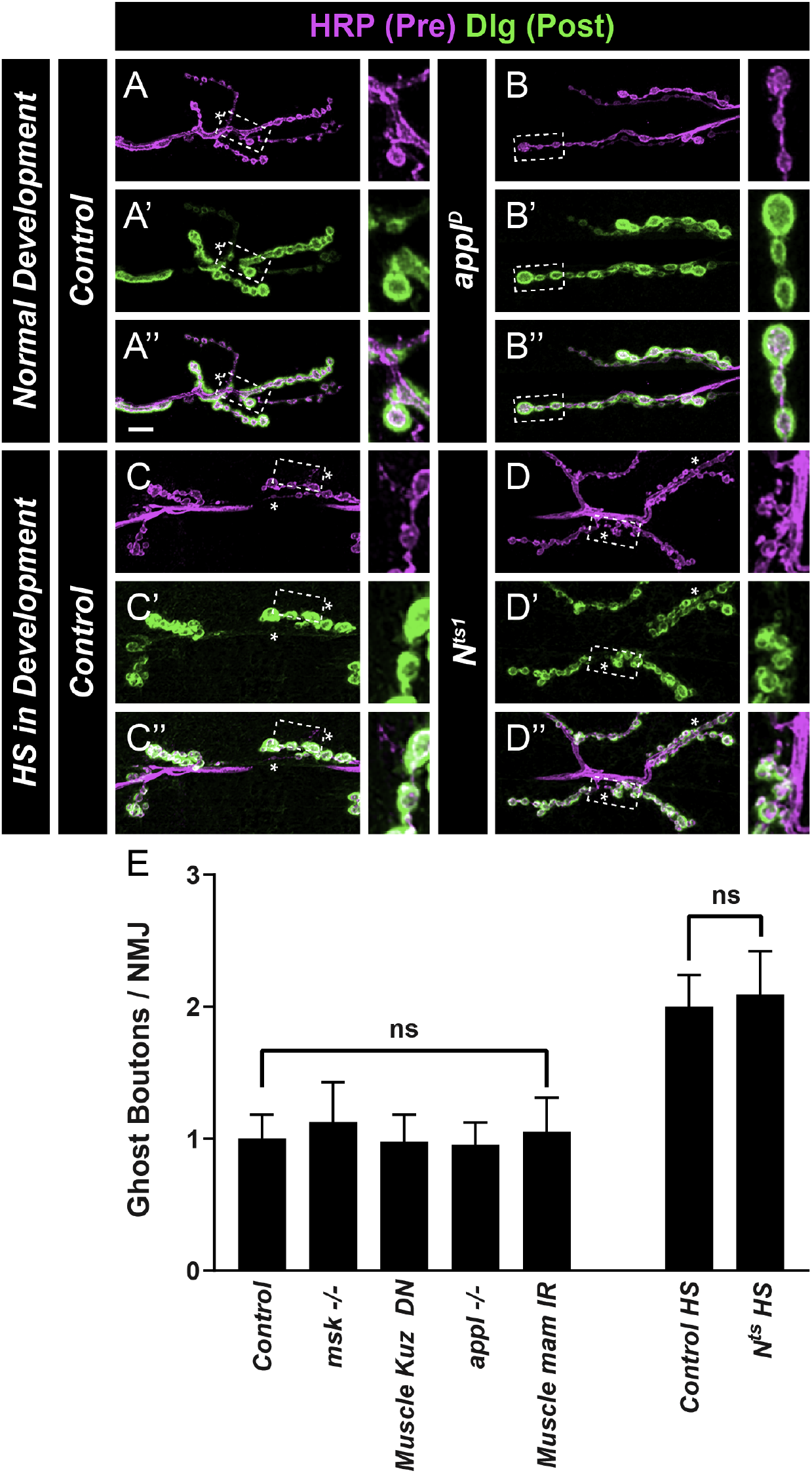
Genetic perturbation of Notch pathway components do not impair postsynaptic development and maturation. (**A – D**) Representative confocal images of NMJs in control (A, C), *appl* mutant (B) and *Notch* mutant (D) larvae and stained with antibodies to HRP (magenta) and Dlg (green). Loss of *appl*, which interacts with the Notch pathway, does not show an increase in ghost boutons. Developmental reduction of Notch itself also does not show an increase in ghost boutons over control larvae. *Control* and *N^TS1^* larvae for this experiment were raised at a higher temperature (29°C) during development, accounting for the increase in ghost boutons when compared to controls raised at 25°C. Insets represent high magnification ghost or control boutons marked by the dashed line. (**E**) Quantification of ghost boutons. In multiple mutants in components of the Notch pathway or muscle-specific perturbation of Notch pathway components, no increases in ghost boutons were observed. This data indicates that perturbing the Notch pathway has no effect on postsynaptic development and maturation, suggesting that γ-secretase does not function through Notch to mediate synaptic maturation. For all *n* ≥ 5 larvae, 10 NMJs. n.s. = not significant. Scale bar = 10 μm.

**Fig. S7.**
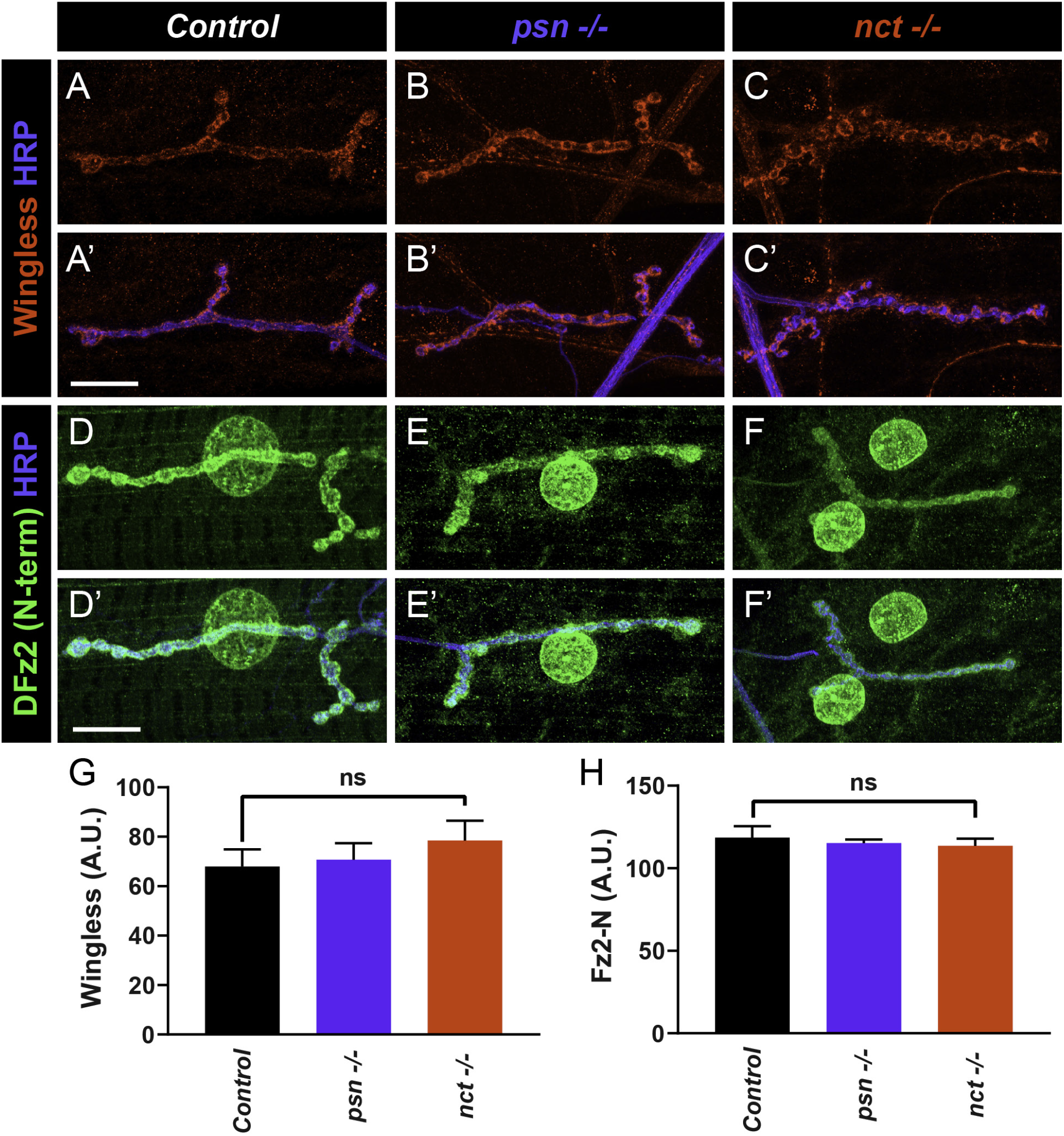
Wnt pathway components are properly localized and trafficked in *psn* and *nct* mutants. (**A – C**) Representative confocal images of control (A), *psn* mutant (B), and *nct* mutant (C) larvae stained with antibodies to the Wnt ligand, Wingless (red) and HRP (blue). (**D – F**) Representative confocal images of control (D), *psn* mutant (E), and *nct* mutant (F) larvae stained with antibodies to the N-terminus of Fz2 (Fz2-N, green) and HRP (blue). Both components properly localize to the NMJ in all mutants observed, suggesting that expression failure cannot account for the observed maturation defects. Fz2-N also decorates the nuclear envelope, further indicating that the endocytosis and trafficking of Fz2 (*14, 23*) is not perturbed in *psn* and *nct* mutants. (**G – H**) Quantification of Wingless (G) and Fz2-N (H) fluorescence. No significant differences in the levels of Wingless or Fz2-N staining are observed in *psn* and *nct* mutants. For all experiments, *n* ≥ 5 larvae, 9 NMJs. n.s. = not significant. Scale bar = 10 μm.

**Fig. S8.**
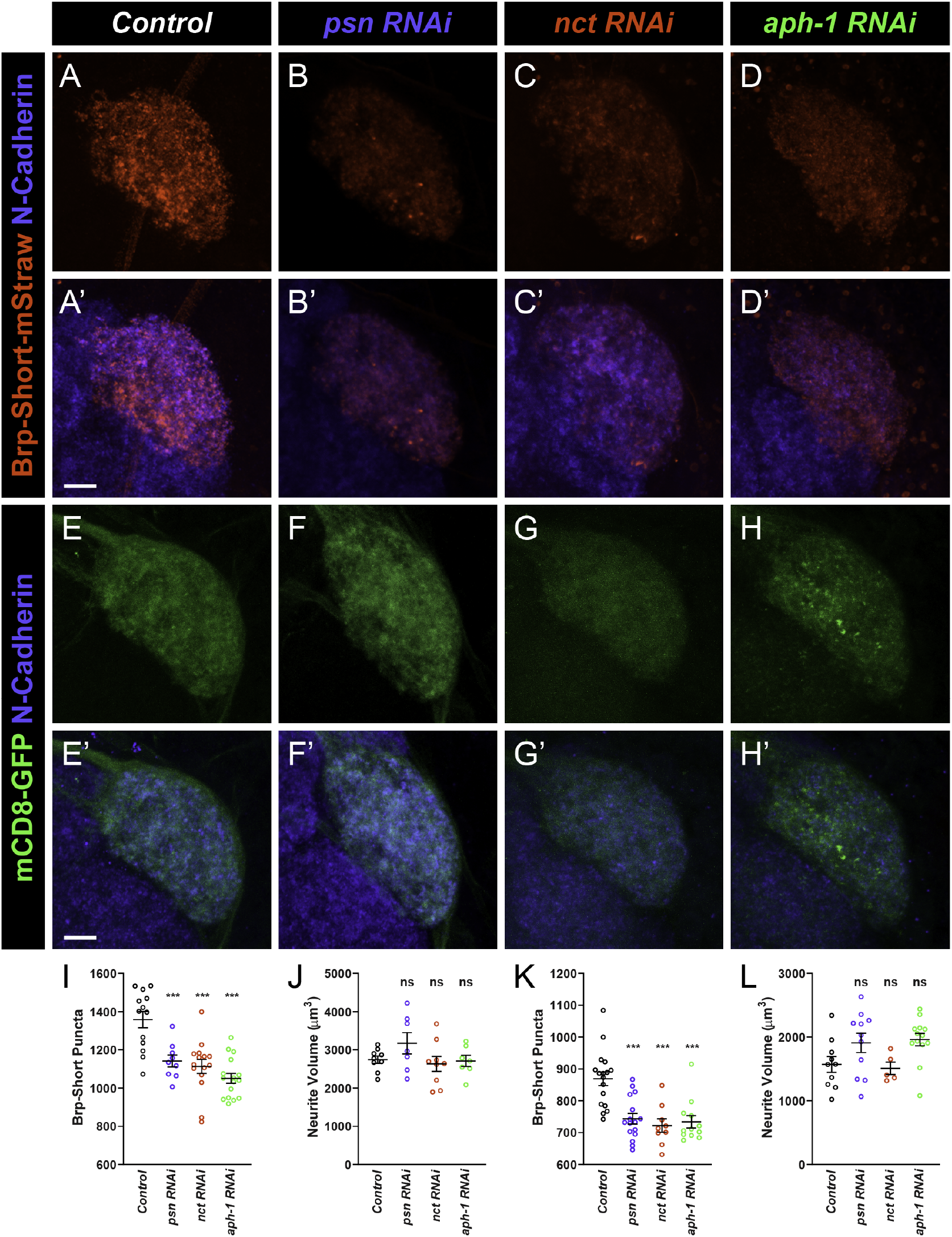
Loss of γ-secretase subunits in olfactory receptor neurons impairs central neuron synaptic development. (**A – D**) Representative high magnification confocal stack images of DA1 olfactory receptor neuron (ORN) axon terminals in the DA1 glomerulus of male adult *Drosophila* expressing Brp-Short-mStraw and stained with antibodies against mStraw (red) and N-Cadherin (blue). RNAi against *psn* (B), *nct* (C), or *aph-1* (D) all show fewer Brp-Short-mStraw puncta compared to control animals (A). (**E – H**) Representative high magnification confocal stack images of DA1 olfactory receptor neuron (ORN) axon terminals in the DA1 glomerulus of male adult *Drosophila* expressing mCD8-GFP and stained with antibodies against GFP (green) and N-Cadherin (blue). RNAi against *psn* (B), *nct* (C), or *aph-1* (D) do not visibly alter GFP staining. (**I, K**) Quantification of Brp-Short-mStraw puncta in DA1 ORNs of male (I) and female (K) flies. There is an 18-27% reduction in Brp-Short-mStraw puncta when γ-secretase subunits are perturbed. (**J, L**) Quantification of mCD8-GFP rendered neurite volume in DA1 ORNs of male (J) and female (L) flies. There are no changes in the neurite volume regardless of genetic perturbation, suggesting that while growth is unaffected, the achievement of a mature number of Brp-Short-mStraw puncta is impaired. This data indicates that γ-secretase in involved in central neuron development in *Drosophila*. For all cases, *n ≥* 5 brains, 9 glomeruli. ***, *p* < 0.001, n.s. = not significant. Scale bar = 20 μm.

**Fig. S9.**
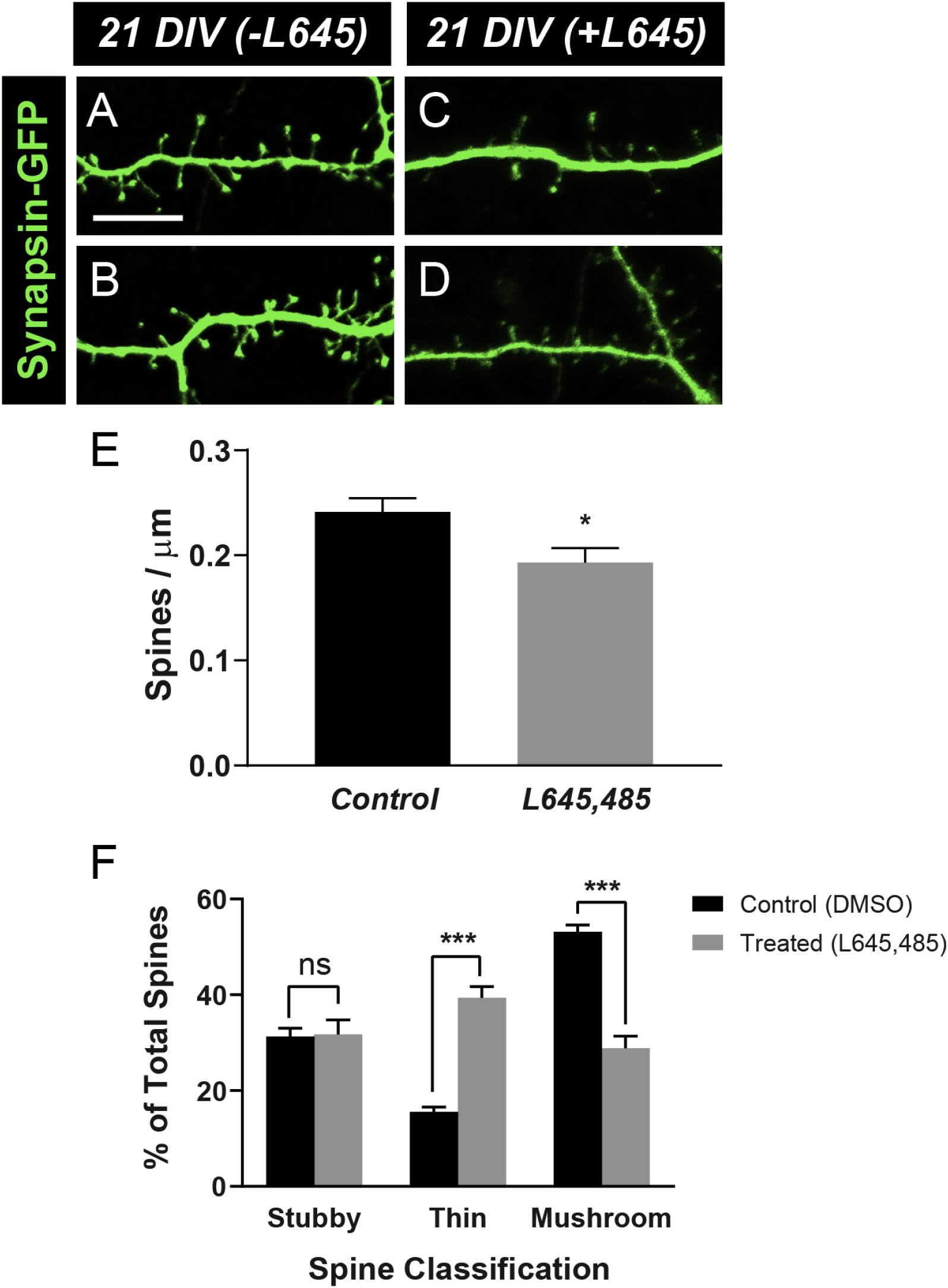
Pharmacologically inhibiting γ-secretase activity impairs dendritic development in primary mammalian neurons. (**A – B**) High magnification of dendritic spines on 21 DIV mammalian primary neuron cultures not treated with L645,485 and stained with antibodies to GFP (green). (**C – D**) 21 DIV cultures stained as above but treated with L645,485. Though dendritic spines are still visible, the mushroom-headed spines in control samples are not often visible; instead, there are more filopodial-like spines. (**E**) Quantification of dendritic spine density. There is a modest reduction in spine density when γ-secretase activity is perturbed, consistent with previous work (*41, 42*). (**F**) Quantification of dendritic spine type. Treated samples show an increase in thin, filopodial outgrowth and a concomitant reduction in mature, mushroom-headed spines. This data indicates that mammalian dendritic development is also impaired by blocking γ-secretase activity, consistent with a conserved role. ***, *p* < 0.001, n.s. = not significant. Scale bar = 10 μm.

