## Supplementary material for "γ-secretase promotes postsynaptic maturation through the cleavage of a Wnt receptor": Table 1

### Table 1: Genotypes for each Figure Panel

| Figure | Panel | Genotype | |
|---|---|---|---|
| 1 | **B** | *+; +; +; +* | |
| | **C** | *w; Mef2-GAL4 / +; UAS-GFP-RNAi / +; +* | |
| | **D** | *w; Mef2-GAL4 / +; UAS-psn-IR-43082 / +; +* | |
| | **E** | *w; Mef2-GAL4 / +; UAS-nct-IR-JF02648 / +; +* | |
| 2 | **A** | *+; +; +; +* | |
| | **B** | *w; +;* $psn^{143/C4}$*; +* | |
| | **C** | *+; +; +; +* | |
| | **D** | *w; +;* $nct^{A7/J2}$*; +* | |
| 3 | **B, F** | *+; +; +; +* | |
| | **C, G** | *w; +;* $psn^{143/C4}$*; +* | |
| | **D, H** | *w; +;* $nct^{A7/J2}$*; +* | |
| | **E, I** | *Control* | *+; +; +; +* |
| | | *psn* | *w; +;* $psn^{143/C4}$*; +* |
| | | *psn + N* | *w; elav-GAL4 / UAS-psn-Nmyc;* $psn^{143/C4}$*; +* |
| | | *psn + M* | *w; Mef2-GAL4 / UAS-psn-Nmyc;* $psn^{143/C4}$*; +* |
| | | *psn RNAi Nerve* | *w; +; elav-GAL4 / UAS-psn-IR-43082; +* |
| | | *psn RNAi Muscle* | *w; Mef2-GAL4 / +; UAS-psn-IR-43082 / +; +* |
| | | *nct* | *w; +;* $nct^{A7/J2}$*; +* |
| | | *nct + N* | *w; elav-GAL4 / +;* $nct^{A7}$ */*$nct^{J2}$*, UAS-nct-myc; +* |
| | | *nct + M* | *w; Mef2-GAL4 / +;* $nct^{A7}$ */*$nct^{J2}$*, UAS-nct-myc; +* |
| | | *nct RNAi Nerve* | *w; +; elav-GAL4 / UAS-nct-IR-JF02648; +* |
| | | *nct RNAi Muscle* | *w; Mef2-GAL4 / +; UAS-nct-IR-JF02648 / +; +* |
| | **K – O** | *Control* | *+; +; +; +* |
| | | *psn* | *w; +;* $psn^{143/C4}$*; +* |
| | | *psn RNAi Nerve* | *w; +; elav-GAL4 / UAS-psn-IR-43082; +* |
| | | *psn RNAi Muscle* | *w; Mef2-GAL4 / +; UAS-psn-IR-43082 / +; +* |
| | | *nct* | *w; +;* $nct^{A7/J2}$*; +* |
| | | *nct RNAi Nerve* | *w; +; elav-GAL4 / UAS-nct-IR-JF02648; +* |
| | | *nct RNAi Muscle* | *w; Mef2-GAL4 / +; UAS-nct-IR-JF02648 / +; +* |
| 4 | **A** | *w; +;* $nct^{J2}$ */ +; +* | |
| | **B** | *w; +;* $dfz2^{C1}$ */ +; +* | |
| | **C** | *w; +;* $nct^{J2}$ */* $dfz2^{C1}$*; +* | |
| | **D** | *Control* | *+; +; +; +* |
| | | *psn -/+* | *w; +;* $psn^{143}$ */ +; +* |
| | | *nct -/+* | *w; +;* $nct^{J2}$ */ +; +* |
| | | *dfz2 -/+* | *w; +;* $dfz2^{C1}$ */ +; +* |
| | | *psn -/+ nct -/+* | *w; +;* $psn^{143}$ */* $nct^{J2}$*; +* |
| | | *psn -/+ dfz2 -/+* | *w; +;* $psn^{143}$ */* $dfz2^{C1}$*; +* |
| | | *nct -/+ dfz2 -/+* | *w; +;* $nct^{J2}$ */* $dfz2^{C1}$*; +* |
| | | *ten-a -/+* | *Df(X)ten-a / +; +; +; +* |
| | | *ten-a -/+ nct -/+* | *Df(X)ten-a / +; +;* $nct^{J2}$ */ +; +* |
| | | *aph-1 -/- dfz2 -/-* | *w;* $aph\text{-}1^{D35/Df}$*;* $dfz2^{C1/Df}$*; +* |
| | **E** | *w; UAS-Psn-Nmyc / UAS-Fz2-FLAG; Mef2-GAL4 / +; +* | |
| | **F** | *+; +; +; +* | |
| | **G** | *+; +; +; +* | |
| | **H** | *w; +; Mef2-GAL4 / UAS-nct-myc; +* | |
| | **I** | *y, w; +; +; +* | |

| | | | |
|---|---|---|---|
| | **J** | y, w; imp-β11$^{70}$ / Df(2R)Δm22; +; + | |
| | **K** | y, w; +; psn$^{143/C4}$; + | |
| | **L** | y, w; +; nct$^{J2/A7}$; + | |
| | **M** | Control | y, w; +; +; + |
| | | psn | y, w; +; psn$^{143/C4}$; + |
| | | psn + N | y, w; elav-GAL4 / UAS-psn-Nmyc; psn$^{143/C4}$; + |
| | | psn + M | y, w; Mef2-GAL4 / UAS-psn-Nmyc; psn$^{143/C4}$; + |
| | | nct | y, w; +; nct$^{A7/J2}$; + |
| | | nct + N | y, w; elav-GAL4 / +; nct$^{A7}$ / nct $^{J2}$, UAS-nct-myc; + |
| | | nct + M | y, w; Mef2-GAL4 / +; nct$^{A7}$ / nct $^{J2}$, UAS-nct-myc; + |
| | **N** | Control | w; Mef2-GAL4 / UAS-Fz2-FLAG; +; + |
| | | β11 -/- | w; imp-β11$^{70}$,UAS-Fz2-FLAG / Df(2R)Δm22; Mef2-GAL4 / +; |
| | | psn -/- | w; Mef2-GAL4 / UAS-Fz2-FLAG; psn$^{143/C4}$; + |
| | | nct -/- | w; Mef2-GAL4 / UAS-Fz2-FLAG; nct$^{A7/J2}$; + |
| 5 | **B, G** | +; +; +; + | |
| | **C, H** | w; Mef2-GAL4 / UAS-GFP-nls; psn$^{143}$; + | |
| | **D, I** | w, UAS-myc-NLS-Fz2C / + or Y; Mef2-GAL4 / +; psn$^{143}$; + | |
| | **E, J** | w; Mef2-GAL4 / UAS-GFP-nls; nct$^{A7/J2}$; + | |
| | **F, K** | w, UAS-myc-NLS-Fz2C / + or Y; Mef2-GAL4 / +; nct$^{A7/J2}$; + | |
| | **L – P** | Control | +; +; +; + |
| | | psn + GFP | w; Mef2-GAL4 / UAS-GFP-nls; psn$^{143}$; + |
| | | psn + Fz2-C | w, UAS-myc-NLS-Fz2C / + or Y; Mef2-GAL4 / +; psn$^{143}$; + |
| | | nct + GFP | w; Mef2-GAL4 / UAS-GFP-nls; nct$^{A7/J2}$; + |
| | | nct + Fz2-C | w, UAS-myc-NLS-Fz2C / + or Y; Mef2-GAL4 / +; nct$^{A7/J2}$; + |
| 6 | **A** | w$^{1118}$; +; +; + | |
| | **B** | w$^{1118}$; +; psn$^{G228D}$; + | |
| | **C** | w; OK371-GAL4 / UAS-Fus$^{WT}$; +; + | |
| | **D** | w; OK371-GAL4 / UAS-Fus$^{R521H}$; +; + | |
| | **E** | w, elav$^{C155}$-GAL4 / + or Y; +; UAS-SNCA-WT / +; + | |
| | **F** | w, elav$^{C155}$-GAL4 / + or Y; +; UAS-SNCA-A53T / +; + | |
| | **G** | w, elav$^{C155}$-GAL4 / + or Y; UAS-HttQ25-GFP / +; +; + | |
| | **H** | w, elav$^{C155}$-GAL4 / + or Y; UAS-HttQ91-GFP / +; +; + | |
| | **I** | Control | w$^{1118}$; +; +; + |
| | | psn$^{G228D}$ | w$^{1118}$; +; psn$^{G228D}$; + |
| | | α-syn-WT | w, elav$^{C155}$-GAL4 / + or Y; +; UAS-SNCA-WT / +; + |
| | | α-syn$^{A30P}$ | w, elav$^{C155}$-GAL4 / + or Y; +; UAS-SNCA-A30P / +; + |
| | | α-syn$^{A53T}$ | w, elav$^{C155}$-GAL4 / + or Y; +; UAS-SNCA-A53T / +; + |
| | | Htt$_{ex1}$Q25 | w, elav$^{C155}$-GAL4 / + or Y; UAS-HttQ25-GFP / +; +; + |
| | | Htt$_{ex1}$Q91 | w, elav$^{C155}$-GAL4 / + or Y; UAS-HttQ91-GFP / +; +; + |
| | | Htt$_{ex1}$Q128 | w, elav$^{C155}$-GAL4 / + or Y; +; UAS-HttQ128 / +; + |
| | | Fus-WT | w; OK371-GAL4 / +; UAS-Fus$^{WT}$ / +; + |
| | | Fus$^{R518K}$ | w, UAS-Fus$^{R518K}$ / + or Y; OK371-GAL4 / +; +; + |
| | | Fus$^{R521C}$ | w; OK371-GAL4 / +; UAS-Fus$^{R521C}$ / +; + |
| | | Fus$^{R521H}$ | w; OK371-GAL4 / UAS-Fus$^{R521H}$; +; + |
| S1 | **B** | +; +; +; + | |
| | **C** | w; +; dfz2$^{C1/Df}$; + | |
| | **D** | w; Mef2-GAL4 / UAS-dshDIX; +; + | |
| | **F** | Control | +; +; +; + |
| | | dfz2 | w; +; dfz2$^{C1/Df}$; + |
| | | dfz2 + FL Fz2 | w; Mef2-GAL4 / UAS-Fz2-FL; dfz2$^{C1/Df}$; + |
| | | dfz2 + Fz2-C | w, UAS-myc-NLS-Fz2C; Mef2-GAL4 / +; dfz2$^{C1/Df}$; + |
| | | dfz2 + ΔKTLES | w; Mef2-GAL4 / UAS-Fz2-DKTLES; dfz2$^{C1/Df}$; + |
| | | Gal4 alone | w; Mef2-GAL4 / +; +; + |

| | | | |
|---|---|---|---|
| | | *Mef2 > Dsh$^{DIX}$* | *w; Mef2-GAL4 / UAS-dshDIX; +; +* |
| S2 | **A – B** | *w; UAS-psn-Nmyc / +; Mef2-GAL4 / +; +* | |
| | **C – D** | *w; +; Mef2-GAL4 / UAS-nct-myc; +* | |
| | **E** | *w; Or47b-GAL4, UAS-Brp-Short-mStraw / UAS-psn-Nmyc; +; +* | |
| | **F** | *w; UAS-Brp-Short-mStraw / +; Or67d-GAL4 / UAS-nct-myc; +* | |
| S3 | **B, F** | *+; +; +; +* | |
| | **C, G** | *w; aph-1$^{D35/Df}$; +; +* | |
| | **D, H** | *w; pen-2$^{MI02639}$; +; +* | |
| | **E,I** | *Control* | *+; +; +; +* |
| | | *aph-1* | *w; aph-1$^{D35/Df}$; +; +* |
| | | *aph-1 + N* | *w; aph-1$^{D35/Df}$; Mef2-GAL4 / UAS-aph-1-V5; +* |
| | | *aph-1 + M* | *w; aph-1$^{D35/Df}$; elav-GAL4 / UAS-aph-1-V5; +* |
| | | *aph-1 RNAi Nerve* | *w; UAS-aph-1-IR-16820 / +; elav-GAL4 / +* |
| | | *aph-1 RNAi Muscle* | *w; UAS-aph-1-IR-16820 / +; Mef2-GAL4 / +; +* |
| | | *pen-2* | *w; pen-2$^{MI02639}$; +; +* |
| | | *pen-2 + N* | *w; pen-2$^{MI02639}$; elav-GAL4 / UAS-pen-2-FLAG; +* |
| | | *pen-2 + M* | *w; pen-2$^{MI02639}$; Mef2-GAL4 / UAS-pen-2-FLAG; +* |
| | | *pen-2 RNAi Nerve* | *w; UAS-pen-2-IR-JF02608 / +; elav-GAL4 / +; +* |
| | | *pen-2 RNAi Muscle* | *w; Mef2-GAL4 / UAS-pen-2-IR-JF02608; +; +* |
| | **K – O** | *Control* | *+; +; +; +* |
| | | *aph-1* | *w; aph-1$^{D35/Df}$; +; +* |
| | | *aph-1 RNAi Nerve* | *w; UAS-aph-1-IR-16820 / +; elav-GAL4 / +* |
| | | *aph-1 RNAi Muscle* | *w; UAS-aph-1-IR-16820 / +; Mef2-GAL4 / +; +* |
| | | *pen-2* | *w; pen-2$^{MI02639}$; +; +* |
| | | *pen-2 RNAi Nerve* | *w; UAS-pen-2-IR-JF02608 / +; elav-GAL4 / +; +* |
| | | *pen-2 RNAi Muscle* | *w; Mef2-GAL4 / UAS-pen-2-IR-JF02608; +; +* |
| S4 | **A,F,K,P** | *+; +; +; +* | |
| | **B,G,L,Q** | *w; +; psn$^{143/C4}$; +* | |
| | **C,H,M,R** | *w; +; nct$^{A7/J2}$; +* | |
| | **D,I,N,S** | *w; aph-1$^{D35/Df}$; +; +* | |
| | **E,J,O,T** | *w; pen-2$^{MI02639}$; +; +* | |
| | **U – Z** | *Control* | *+; +; +; +* |
| | | *psn* | *w; +; psn$^{143/C4}$; +* |
| | | *nct* | *w; +; nct$^{A7/J2}$; +* |
| | | *aph-1* | *w; aph-1$^{D35/Df}$; +; +* |
| | | *pen-2* | *w; pen-2$^{MI02639}$; +; +* |
| S5 | **A – B** | *+; +; +; +* | |
| | **C** | *Control* | *+; +; +; +* |
| | | *Psn* | *w; +; psn$^{143/C4}$; +* |
| S6 | **A** | *+; +; +; +* | |
| | **B** | *w, Appl$^{d}$; +; +; +* | |
| | **C** | *+; +; +; +* | |
| | **D** | *N$^{ts1}$; +; +; +* | |
| | **E** | *Control* | *+; +; +; +* |
| | | *Msk -/-* | *w; +; msk$^{5}$; +* |
| | | *Muscle Kuz DN* | *w; UAS-Kuz-DN / +; Mef2-GAL4 / +; +* |
| | | *Appl -/-* | *w, Appl$^{d}$; +; +; +* |
| | | *Muscle mam IR* | *w; Mef2-GAL4 / UAS-mam-IR-102091; +; +* |
| | | *Control HS* | *+; +; +; +* |
| | | *N$^{ts}$ HS* | *N$^{ts1}$; +; +; +* |
| S7 | **A,D** | *+; +; +; +* | |
| | **B,E** | *w; +; psn$^{143/C4}$; +* | |

| | | | |
|---|---|---|---|
| | C,F | *w; +; $nct^{A7/J2}$; +* | |
| | G,H | *Control* | *+; +; +; +* |
| | | *psn -/-* | *w; +; $psn^{143/C4}$; +* |
| | | *nct -/-* | *w; +; $nct^{A7/J2}$; +* |
| S8 | A | *w, UAS-Dcr2 / Y; UAS-Brp-Short-Straw / +; Or67d-GAL4 / +; +* | |
| | B | *w, UAS-Dcr2 / Y; UAS-Brp-Short-Straw / +; Or67d-GAL4 / UAS-psn-IR-43082; +* | |
| | C | *w, UAS-Dcr2 / Y; UAS-Brp-Short-Straw / +; Or67d-GAL4 / UAS-nct-IR-JF02648; +* | |
| | D | *w, UAS-Dcr2 / Y; UAS-Brp-Short-Straw / UAS-aph-1-IR-16820; Or67d-GAL4 / +; +* | |
| | E | *w, UAS-Dcr2 / Y; +; Or67d-GAL4, UAS-mCD8-GFP / +; +* | |
| | F | *w, UAS-Dcr2 / Y; +; Or67d-GAL4, UAS-mCD8-GFP / UAS-psn-IR-43082; +* | |
| | G | *w, UAS-Dcr2 / Y; +; Or67d-GAL4, UAS-mCD8-GFP / UAS-nct-IR-JF02648; +* | |
| | H | *w, UAS-Dcr2 / Y; UAS-aph-1-IR-16820 / +; Or67d-GAL4, UAS-mCD8-GFP / +; +* | |
| | I | *Control* | *w, UAS-Dcr2 / Y; UAS-Brp-Short-Straw / +; Or67d-GAL4 / +; +* |
| | | *psn RNAi* | *w, UAS-Dcr2 / Y; UAS-Brp-Short-Straw / +; Or67d-GAL4 / UAS-psn-IR-43082; +* |
| | | *nct RNAi* | *w, UAS-Dcr2 / Y; UAS-Brp-Short-Straw / +; Or67d-GAL4 / UAS-nct-IR-JF02648; +* |
| | | *aph-1 RNAi* | *w, UAS-Dcr2 / Y; UAS-Brp-Short-Straw / UAS-aph-1-IR-16820; Or67d-GAL4 / +; +* |
| | J | *Control* | *w, UAS-Dcr2 / w; UAS-Brp-Short-Straw / +; Or67d-GAL4 / +; +* |
| | | *psn RNAi* | *w, UAS-Dcr2 / w; UAS-Brp-Short-Straw / +; Or67d-GAL4 / UAS-psn-IR-43082; +* |
| | | *nct RNAi* | *w, UAS-Dcr2 / w; UAS-Brp-Short-Straw / +; Or67d-GAL4 / UAS-nct-IR-JF02648; +* |
| | | *aph-1 RNAi* | *w, UAS-Dcr2 / Y; UAS-Brp-Short-Straw / UAS-aph-1-IR-16820; Or67d-GAL4 / +; +* |
| | K | *Control* | *w, UAS-Dcr2 / w; +; Or67d-GAL4, UAS-mCD8-GFP / +; +* |
| | | *psn RNAi* | *w, UAS-Dcr2 / Y; +; Or67d-GAL4, UAS-mCD8-GFP / UAS-psn-IR-43082; +* |
| | | *nct RNAi* | *w, UAS-Dcr2 / Y; +; Or67d-GAL4, UAS-mCD8-GFP / UAS-nct-IR-JF02648; +* |
| | | *aph-1 RNAi* | *w, UAS-Dcr2 / Y; UAS-aph-1-IR-16820 / +; Or67d-GAL4, UAS-mCD8-GFP / +; +* |
| | L | *Control* | *w, UAS-Dcr2 / w; +; Or67d-GAL4, UAS-mCD8-GFP / +; +* |
| | | *psn RNAi* | *w, UAS-Dcr2 / w; +; Or67d-GAL4, UAS-mCD8-GFP / UAS-psn-IR-43082; +* |
| | | *nct RNAi* | *w, UAS-Dcr2 / w; +; Or67d-GAL4, UAS-mCD8-GFP / UAS-nct-IR-JF02648; +* |
| | | *aph-1 RNAi* | *w, UAS-Dcr2 / w; UAS-aph-1-IR-16820 / +; Or67d-GAL4, UAS-mCD8-GFP / +; +* |
